# To pack or not to pack: revisiting protein side-chain packing in the post-AlphaFold era

**DOI:** 10.1101/2025.02.22.639681

**Authors:** Sriniketh Vangaru, Debswapna Bhattacharya

## Abstract

**Motivation:** Protein side-chain packing (PSCP), the problem of predicting side-chain conformation given a fixed backbone structure, has important implications in modeling of structures and interactions. However, despite the groundbreaking progress in protein structure prediction pioneered by AlphaFold, the existing PSCP methods still rely on experimental inputs, and do not leverage AlphaFold-predicted backbone coordinates to enable PSCP at scale.

**Results:** Here, we perform a large-scale benchmarking of the predictive performance of various PSCP methods on public datasets from multiple rounds of the Critical Assessment of Structure Prediction (CASP) challenges using a diverse set of evaluation metrics. Empirical results demonstrate that the PSCP methods perform well in packing the side-chains with experimental inputs, but they fail to generalize in repacking AlphaFold-generated structures. We additionally explore the effectiveness of leveraging the self-assessment confidence scores from AlphaFold by implementing a backbone confidence-aware integrative approach. While such a protocol often leads to performance improvement by attaining modest yet statistically significant accuracy gains over the AlphaFold baseline, it does not yield consistent and pronounced improvements. Our study highlights the recent advances and remaining challenges in PSCP in the post-AlphaFold era.

**Availability:** The code and raw data are freely available at https://github.com/Bhattacharya-Lab/PackBench.

## 1. Introduction

Predicting the precise three-dimensional (3D) configuration of the side-chain atoms of a protein molecule given the arrangement of its backbone atoms, the so-called protein side-chain packing (PSCP) problem, is critically important for high-accuracy modeling of macromolecular structures [1– 7] and interactions [8–13]. Over the past several decades, the PSCP task has gained significant attention and garnered considerable research efforts [14, 15]. The existing PSCP methods can be broadly categorized into three major classes: rotamer (rotational isomer) library-based algorithms [16–25], probabilistic or machine learning-based approaches [26, 27], and deep learning- or generative modeling-based methods [28–36]. Side-chain predictors that rely on rotamer libraries generally focus on minimizing the global energy of the protein structure using stochastic optimization [19, 20, 24] and/or graph theoretic algorithms [23, 25]. Beyond rotamers, probabilistic or machine learning-based approaches have been developed which implicitly model the side-chain conformational space using a dynamic Bayesian network [27] or combine neural networks with Markov Chain Monte Carlo optimization through a hybrid approach [26]. More recently, a number of deep learning-based methods have shown promising results by making use of various representations and deep neural network frameworks, ranging from estimating side-chain densities with a voxelized representation of each residue’s local environment using a U-net-style architecture [32] to directly predicting the side-chain coordinates with attention-based transformer networks [33]to predicting side-chain torsion angle distributions using a geometry-aware invariant point message passing architecture [35]. Some of the emerging deep generative models such as a torsional diffusion model for autoregressive side-chain packing or torsional flow matching [36] represent the state of the art in PSCP, achieving impressive accuracy when experimentally resolved backbone coordinates are used as input.

Alongside these developments in PSCP, recent breakthroughs in sequence-based protein structure prediction by AlphaFold2 [37] and AlphaFold3 [38] have revolutionized structural biology [39] by enabling highly accurate prediction of protein structures that can approach near-experimental quality [40] while achieving massive scalability compared to experimental structure determination methods [41]. Notably, the AlphaFold methods not only enable all-atom prediction of 3D protein structures containing both backbone and side-chain atoms, but also provides self-assessment confidence scores by estimating the accuracy of their own predictions even in the absence of the native (experimental) structure. Despite such advances, the vast majority of the existing PSCP methods still primarily rely on experimental backbone as their input as opposed to leveraging AlphaFold-predicted backbone coordinates. Given the recent progress, a natural question arises: how robust are the existing PSCP methods in leveraging the backbone conformations predicted by AlphaFold as inputs for side-chain packing, in order to further improve side-chain positioning beyond what is possible by AlphaFold?

Here, we benchmark the predictive performance of a representative collection of the existing PSCP methods over a large set of protein targets from the 14^th^ and 15^th^ rounds of the Critical Assessment of Structure Prediction (CASP) challenges using various complementary performance evaluation metrics. Our objective and empirical study has three major contributions. First, recognizing that the existing PSCP methods traditionally rely on experimental backbone coordinates as inputs, we present a performance benchmark based on the native backbones. Second, we assess the ability of the PSCP methods in repacking the side-chain conformations when AlphaFold2- and AlphaFold3-generated backbone coordinates are used as inputs, and investigate to what extent such repacking improves the baseline side-chain accuracy achieved by AlphaFold. Third, we explore the possibility of leveraging the self-assessment confidence scores from AlphaFold by implementing an integrative approach that combines a variety of PSCP methods to further improve the fidelity of side-chain prediction beyond the baseline performance of AlphaFold. The code and raw data are freely available at https://github.com/Bhattacharya-Lab/PackBench.

## 2. Methods

### 2.1. Benchmarking datasets

We used protein targets having length less than 2,000 residues from the 14^th^ and 15^th^ editions of the CASP experiments as our benchmarking datasets. The CASP14 set contained 66 single-chain targets, while the CASP15 set consisted of 71 single-chain targets. Further details on the dataset preparation can be found in **Section 1.1** of the **Supplementary Information**. For each of these targets, we collected structures predicted by AlphaFold2 and AlphaFold3 based on the CASP target sequences [37, 38]. AlphaFold2’s predictions of the CASP14 targets were retrieved directly from the CASP14 data archive (https://predictioncenter.org/download_area/CASP14/predictions/regular/), while its predictions of the CASP15 targets were downloaded from AlphaFold2’s public release of CASP15 structures on GitHub at https://github.com/google-deepmind/alphafold/blob/main/docs/technical_note_v2.3.0.md. We generated all AlphaFold3 predictions by submitting the sequences into the AlphaFold3 online server available at https://alphafoldserver.com/.

### 2.2. Side-chain packing methods assessed

We evaluated a diverse set of PSCP methods for our assessment. First, we ran each method with the native backbone coordinates in order to benchmark their performance on their expected inputs. Subsequently, we replaced the native backbones with AlphaFold-predicted backbone coordinates as inputs (for both CASPs and both AlphaFold versions) to investigate the impact of this change. Further details on how we installed and ran inference using the PSCP methods are provided in **Section 1.2** of our **Supplementary Information**. Below, we briefly summarize the PSCP methods assessed.

1. *SCWRL4:* SCWRL4 [23] is one of the most widely used rotamer library-based PSCP algorithms that leverages backbone-dependent rotamer conformations [42].
2. *Rosetta Packer:* Rosetta Packer, a part of the Rosetta3 software suite [24], performs PSCP using a rotamer library through Rosetta energy minimization. In our study, we use the PyRosetta package [43], which implements the core Rosetta functionality in Python.
3. *FASPR:* FASPR [25] uses an optimized scoring function in combination with a deterministic search algorithm for rotamer library-based PSCP.
4. *DLPacker:* DLPacker [32], one of the first deep learning-based PSCP methods, is based on a voxelized representation of each residue’s local environment using a U-net-style architecture.
5. *AttnPacker:* AttnPacker [33] is an end-to-end, SE(3)-equivariant deep graph transformer architecture for the direct prediction of side-chain coordinates, and it additionally implements a post-processing procedure in order to reduce steric clashes, aiming to produce physically realistic conformations.
6. *DiffPack:* DiffPack [34] leverages a torsional diffusion model for PSCP in an autoregressive manner, progressively conditioning each *χ* angle on its output for the previous angles.
7. *PIPPack:* PIPPack [35], also known as the Protein Invariant Point Packer, processes local structural and sequence information for PSCP using *χ*-angle distribution predictions and invariant point message passing (IPMP), a generalization of the invariant point attention (IPA) module introduced in AlphaFold2 [37].
8. *FlowPacker:* FlowPacker [36] uses torsional flow matching for PSCP by training a continuous normalizing flow (CNF) model with equivariant graph attention networks.

### 2.3. Repacking AlphaFold side-chains using a backbone confidence-aware integrative approach

AlphaFold provides self-assessment confidence scores by estimating the Local Distance Difference Test (lDDT) [44] in the form of their so-called predicted lDDT (plDDT)— with residue-level granularity for AlphaFold2 and atom-level granularity for AlphaFold3—which ranges from 0 to 100, quantifying how confident AlphaFold is in its own prediction. We investigate the possibility of augmenting AlphaFold’s baseline side-chain performance by integrating this self-assessment capability with various PSCP methods. Specifically, our approach to repack an AlphaFold-generated protein structure’s side-chains acts as a post-processing step for AlphaFold’s pipeline, using a greedy energy minimization scheme to search for more optimal *χ* angles in the rotamer conformation space. It begins by initializing a structure equal to AlphaFold’s output, and generates variations of the structure by using each of the existing tools to repack the side-chains of this structure. We then search for improved *χ* angles by greedily minimizing the 2015 Rosetta Energy Function (REF2015) [45]—which has been shown to be effective in capturing the all-atom protein conformational space [46] and has been used widely in protein modeling and refinement [47– 49]. The algorithm repeatedly selects a *χ* angle *j* from a residue *i* and a tool *k*, updating the angle in the current structure to a weighted average of itself and the corresponding angle from tool *k*’s prediction only if that operation lowers the energy of the overall structure. We integrate residue *i*’s backbone plDDT into this step of the algorithm by using it as the weight of the current structure’s *χ* angle, which effectively biases the search process to stick closer to more confident AlphaFold predictions (see **Algorithm 1** in **Supplementary Information** for more details). It is worth noting that such a weighing scheme works for both AlphaFold2 and AlphaFold3: AlphaFold2 provides residue-wise plDDT values, which are treated as backbone self-assessment confidence scores, whereas the per-atom plDDT scores provided by AlphaFold3 are averaged over the backbone heavy atoms to produce the backbone confidence scores at the residue level.

### 2.4. Evaluation metrics and performance assessment

We used standard evaluation metrics for performance benchmarking of the various PSCP methods. First, we calculated the root mean square deviation (RMSD) to quantify the distances between corresponding atoms in the predicted and native structures based on optimal structural superposition. This was separately calculated across all amino acid residues, residues in the core of the protein, and residues at the surface. By defining the *centrality* of a residue to be the number of other residues’ *C*_*β*_ atoms within a 10°A radius of that residue’s *C*_*β*_ atom, *core* residues are those with centrality ≥ 20 and *surface* residues are those with centrality ≤ 15. Second, the mean absolute error (MAE) of the *χ*_1_ – *χ*_4_ torsion angles of each residue was used for determining the accuracy of side-chains in the torsional space. A closely related evaluation measure is the recovery rate (RR), measured as the percentage of residues for which all 4 *χ* angles have an MAE under 20°. Lastly, in order to measure physical realism, we compute the total count of steric clashes per target, where a *clash* is defined as a pair of atoms which are not bonded and yet whose pairwise distance is less than a proportion of the sum of their van der Waals radii (the thresholds of proportions used were 100%, 90%, and 80%).

To determine if any of these PSCP methods demonstrated a noticeable improvement over the AlphaFold baseline structures that they repacked, we additionally performed statistical significance tests that analyzed the degree of this change. For each combination of original dataset (CASP14 or CASP15), predicted backbone generated by AlphaFold2 or AlphaFold3, PSCP method used for repacking the side-chains, and metric being evaluated, a hypothesis test was ran using all of the *n* ∈ {66, 71} predicted structures at the intersection of those selections, resulting in 792 total statistical tests. We used Wilcoxon signed-rank tests to avoid any assumptions about the parameters of the respective distributions of metric values, and each individual before-after pairing consisted of the original AlphaFold-predicted side-chains for a given target paired with the repacked side-chains for that target. Each test was considered as one-tailed using *α* = 0.05, with the ‘null’ group representing no statistically significant improvement, or a decrease, in performance on the corresponding metric. An improvement is defined to be an increase for the recovery rate of the structure and a decrease for every other metric. Statistical tests in this same form were conducted for our backbone confidence-aware integrative approach to evaluate its effectiveness in repacking AlphaFold’s side-chains beyond its baseline accuracy.

## 3. Results and discussion

### 3.1. Performance benchmarking with experimental backbone coordinates as input

**Table 1** presents the performance benchmarking with experimental backbone coordinates as input. The benchmarking reveals some interesting insights. First, the RMSD tends to be lower for the core residues compared to that of the surface residues, likely due to less conformational flexibility of the core residues [50], in part due to the hydrophobic packing effect [51]. Second, *χ*_3_ and *χ*_4_ show noticeably higher mean absolute errors in comparison to that of *χ*_1_ and *χ*_2_, possibly due to their high side-chain torsional degrees of freedom. Third, across the various metrics evaluated, the deep learning-based methods consistently outperformed the traditional PSCP methods. One exception to this is the PyRosetta Packer which attained the best results for steric clashes, likely due to the use of the biophysics-informed REF15 energy function. Among the deep learning-based methods, the three most recent ones (FlowPacker, PIPPack, DiffPack) had substantially higher rotamer recovery rates than all other tools on both CASP sets, demonstrating the effectiveness of deep learning and generative modeling approaches for PSCP.

**Table 1.** Performance benchmarking of the PSCP methods on CASP targets with native backbone coordinates as inputs.

| Tool Name | RMSD (Å) ↓ | | | χ-MAE (°) ↓ | | | | RR (%) ↑ | Steric Clashes (#) ↓ | | |
| --- | --- | --- | --- | --- | --- | --- | --- | --- | --- | --- | --- |
| | All | Core | Surface | $\chi_1$ | $\chi_2$ | $\chi_3$ | $\chi_4$ | $\chi_{1-4}$ | 100% | 90% | 80% |
| **CASP14** ($n = 66$) | | | | | | | | | | | |
| FlowPacker<sup>1</sup> | 0.80 | **0.40** | 1.01 | 23.02 | 25.82 | 46.09 | 52.80 | 57.1 | 102.0 | 21.7 | 6.4 |
| PIPPack<sup>2</sup> | 0.79 | 0.43 | 0.99 | **21.57** | 25.25 | **41.93** | 51.27 | **58.1** | 131.2 | 36.2 | 14.4 |
| DiffPack<sup>3</sup> | **0.79** | 0.41 | 0.98 | 22.92 | **25.23** | 46.97 | 55.33 | 57.6 | 104.2 | 26.8 | 9.8 |
| AttnPacker<sup>4</sup> | 0.79 | 0.44 | **0.98** | 24.19 | 28.79 | 48.34 | **50.37** | 51.3 | 84.6 | 22.8 | 8.1 |
| DLPacker<sup>5</sup> | 0.90 | 0.50 | 1.11 | 27.45 | 30.03 | 52.82 | 70.34 | 50.6 | **83.2** | **16.8** | **5.1** |
| FASPR | 1.03 | 0.62 | 1.24 | 31.97 | 31.27 | 49.43 | 55.74 | 47.8 | 152.9 | 41.8 | 13.0 |
| PyRosetta Packer | 1.00 | 0.55 | 1.23 | 30.98 | 31.29 | 49.31 | 55.58 | 48.9 | 104.3 | 22.1 | 8.4 |
| SCWRL4 | 1.04 | 0.61 | 1.26 | 32.22 | 31.65 | 50.21 | 55.10 | 47.5 | 158.3 | 40.2 | 11.8 |
| **CASP15** ($n = 71$) | | | | | | | | | | | |
| FlowPacker | 0.69 | **0.33** | 0.90 | 18.99 | **22.04** | 40.93 | **52.62** | **66.4** | 100.8 | 14.6 | 3.3 |
| PIPPack | 0.70 | 0.34 | 0.91 | **18.27** | 22.16 | **40.21** | 53.36 | 66.1 | 129.0 | 30.5 | 10.9 |
| DiffPack | **0.68** | 0.34 | **0.87** | 18.29 | 22.47 | 42.91 | 56.88 | 65.7 | 95.3 | 20.3 | 7.2 |
| AttnPacker | 0.71 | 0.37 | 0.90 | 20.29 | 26.10 | 47.09 | 54.68 | 59.2 | 96.4 | 25.5 | 9.5 |
| DLPacker | 0.76 | 0.38 | 0.97 | 21.88 | 26.29 | 50.86 | 67.53 | 59.5 | **89.4** | 14.0 | 3.2 |
| FASPR | 0.92 | 0.52 | 1.14 | 27.12 | 29.07 | 50.39 | 59.05 | 55.8 | 160.5 | 37.4 | 9.7 |
| PyRosetta Packer | 0.87 | 0.43 | 1.12 | 25.84 | 27.57 | 47.95 | 55.32 | 58.0 | 98.5 | **13.5** | **3.1** |
| SCWRL4 | 0.94 | 0.50 | 1.17 | 27.89 | 29.12 | 49.81 | 57.25 | 55.5 | 168.3 | 36.3 | 7.7 |

Values in bold indicate the best performance.
<sup>1</sup>“FlowPacker” refers to the variation trained on a snapshot of the Protein Data Bank (PDB) clustered using 40% sequence identity and using confidence sampling with 8 samples.
<sup>2</sup>“PIPPack” refers to the variation with ensembled inferences.
<sup>3</sup>“DiffPack” refers to the variation with confidence sampling.
<sup>4</sup>“AttnPacker” refers to the variation without any postprocessing for steric clashes.
<sup>5</sup>“DLPacker” refers to the variation that passes through residues in an order based on the scores of predictions from a separate pre-pass.
Performance of all other variants of the PSCP methods are found in **Section 2** of our **Supplementary Information**.

### 3.2. Performance benchmarking with AlphaFold2 and AlphaFold3 backbone coordinates as input

To evaluate the effectiveness of the PSCP methods in the post-AlphaFold era, in **Table 2** we report their performance in repacking the side-chain conformations when AlphaFold2- and AlphaFold3-generated backbone coordinates are used as inputs instead of experimental inputs, alongside the performance of the side-chains predicted by the AlphaFold methods as a baseline. Interestingly, the baseline AlphaFold-generated side-chains exhibit the best performance for almost all metrics compared to any of the PSCP methods in repacking AlphaFold-generated backbones. **Figure 1** shows that across the various metrics used, the relative changes attained by the PSCP methods in repacking AlphaFold side-chains from the AlphaFold baseline performance lean toward the undesirable direction, indicating degradation in side-chain accuracy compared to AlphaFold. The Wilcoxon signed-rank tests reveal that the performance improvement due to repacking by the PSCP methods, if any, are statistically indistinguishable from the ‘null’ group representing the baseline AlphaFold (see **Supplementary Tables 2, 3, 4, 5**). It is interesting to note, however, that the performance of the PSCP methods with experimental backbone coordinates as their input reported in **Table 1** are consistently better than AlphaFold’s baseline side-chain performance. That is, PSCP methods have significantly lower performance in repacking the side-chain conformations when AlphaFold2- and AlphaFold3-generated backbone coordinates are used as inputs than when they are given experimental inputs. Such a lack of generalizability puts into question the practical relevance of the existing PSCP methods in the post-AlphaFold era where proteome-wide structures can be predicted by AlphaFold methods [41].

**Table 2.** Performance benchmarking of the PSCP methods on CASP targets with AlphaFold2 and AlphaFold3 backbone coordinates as inputs.

| Tool Name | RMSD (Å) ↓ | | | χ-MAE (°) ↓ | | | | RR (%) ↑ | Steric Clashes (#) ↓ | | |
| --- | --- | --- | --- | --- | --- | --- | --- | --- | --- | --- | --- |
| | All | Core | Surface | $\chi_1$ | $\chi_2$ | $\chi_3$ | $\chi_4$ | $\chi_{1-4}$ | 100% | 90% | 80% |
| **CASP14** ($n = 66$) | | | | | | | | | | | |
| **AF2-Generated Backbones** |  |  |  |  |  |  |  |  |  |  |  |
| AlphaFold2 | 1.07 | **0.66** | 1.26 | **34.59** | **31.51** | **50.81** | **51.42** | 46.0 | **41.4** | **1.9** | **0.0** |
| FlowPacker | 1.09 | 0.67 | 1.30 | 35.62 | 33.04 | 51.12 | 55.85 | **46.1** | 86.1 | 13.1 | 2.7 |
| PIPPack | 1.10 | 0.68 | 1.30 | 35.71 | 33.06 | 51.07 | 54.55 | 45.1 | 102.3 | 20.6 | 6.5 |
| DiffPack | 1.12 | 0.70 | 1.34 | 36.69 | 32.99 | 53.41 | 56.13 | 44.9 | 57.2 | 11.7 | 3.7 |
| AttnPacker | **1.06** | 0.68 | **1.25** | 36.08 | 34.92 | 52.85 | 51.78 | 43.3 | 68.5 | 15.3 | 4.5 |
| DLPacker | 1.11 | 0.70 | 1.33 | 36.28 | 35.19 | 57.87 | 72.25 | 42.9 | 67.7 | 11.0 | 2.1 |
| FASPR | 1.18 | 0.75 | 1.38 | 38.94 | 34.70 | 53.43 | 55.79 | 41.9 | 121.1 | 27.0 | 5.7 |
| PyRosetta Packer | 1.16 | 0.72 | 1.38 | 38.16 | 35.19 | 52.84 | 55.31 | 42.7 | 73.9 | 7.7 | 1.2 |
| SCWRL4 | 1.20 | 0.77 | 1.40 | 39.01 | 35.09 | 52.93 | 56.20 | 41.6 | 132.8 | 29.0 | 5.7 |
| **AF3-Generated Backbones** |  |  |  |  |  |  |  |  |  |  |  |
| AlphaFold3 | **1.04** | **0.64** | 1.25 | **34.14** | **30.35** | **49.42** | **50.31** | **47.4** | **45.8** | **5.2** | **0.7** |
| FlowPacker | 1.07 | 0.66 | 1.29 | 34.85 | 31.91 | 51.16 | 54.77 | 47.3 | 79.0 | 11.6 | 2.5 |
| PIPPack | 1.08 | 0.67 | 1.29 | 35.18 | 32.38 | 50.02 | 52.42 | 46.6 | 95.1 | 19.5 | 6.3 |
| DiffPack | 1.11 | 0.69 | 1.33 | 36.43 | 32.66 | 51.65 | 56.08 | 45.8 | 56.0 | 12.1 | 4.0 |
| AttnPacker | 1.04 | 0.66 | **1.24\*** | 35.26 | 34.11 | 51.57 | 51.90 | 43.9 | 62.4 | 14.3 | 4.3 |
| DLPacker | 1.10 | 0.69 | 1.31 | 36.28 | 34.13 | 56.91 | 72.26 | 43.3 | 64.5 | 10.3 | 2.2 |
| FASPR | 1.16 | 0.76 | 1.36 | 38.06 | 33.48 | 52.29 | 53.81 | 43.2 | 115.3 | 25.6 | 5.7 |
| PyRosetta Packer | 1.15 | 0.71 | 1.37 | 37.66 | 34.51 | 52.23 | 54.16 | 43.4 | 69.3 | 7.1 | 1.4 |
| SCWRL4 | 1.17 | 0.77 | 1.37 | 37.69 | 34.12 | 51.83 | 57.18 | 43.1 | 127.7 | 28.1 | 6.4 |
| **CASP15** ($n = 71$) | | | | | | | | | | | |
| **AF2-Generated Backbones** |  |  |  |  |  |  |  |  |  |  |  |
| AlphaFold2 | **0.90** | **0.58** | **1.11** | **28.05** | **27.90** | **48.04** | **55.00** | 53.9 | **48.2** | **2.0** | **0.0** |
| FlowPacker | 0.94 | 0.59 | 1.15 | 29.38 | 29.18 | 50.18 | 57.00 | **55.1\*** | 100.4 | 13.9 | 2.3 |
| PIPPack | 0.96 | 0.61 | 1.19 | 30.08 | 29.89 | 50.22 | 56.32 | 53.5 | 124.5 | 27.3 | 9.4 |
| DiffPack | 0.99 | 0.63 | 1.21 | 31.07 | 30.10 | 51.90 | 56.39 | 52.8 | 69.7 | 13.7 | 4.5 |
| AttnPacker | 0.92 | 0.61 | 1.12 | 30.34 | 31.48 | 52.10 | 55.52 | 50.4 | 87.9 | 23.6 | 7.7 |
| DLPacker | 0.97 | 0.62 | 1.18 | 30.94 | 31.05 | 55.90 | 69.34 | 51.3 | 84.2 | 12.9 | 2.7 |
| FASPR | 1.06 | 0.71 | 1.27 | 33.67 | 32.53 | 53.56 | 59.75 | 50.0 | 147.0 | 32.5 | 7.9 |
| PyRosetta Packer | 1.04 | 0.67 | 1.26 | 32.78 | 32.67 | 51.42 | 58.65 | 51.0 | 91.6 | 10.7 | 2.4 |
| SCWRL4 | 1.07 | 0.71 | 1.29 | 33.95 | 32.73 | 53.35 | 61.55 | 50.0 | 160.7 | 35.0 | 7.5 |
| **AF3-Generated Backbones** |  |  |  |  |  |  |  |  |  |  |  |
| AlphaFold3 | **0.95** | **0.60** | 1.16 | **30.18** | **28.90** | **48.92** | **53.94** | **53.8** | **58.4** | **8.1** | **1.0** |
| FlowPacker | 0.98 | 0.62 | 1.19 | 31.12 | 30.44 | 50.88 | 57.02 | 53.7 | 92.5 | 13.3 | 2.5 |
| PIPPack | 1.00 | 0.64 | 1.22 | 31.57 | 30.65 | 49.81 | 57.75 | 52.7 | 115.7 | 26.5 | 9.0 |
| DiffPack | 1.02 | 0.66 | 1.23 | 32.44 | 31.03 | 52.27 | 61.20 | 51.6 | 70.8 | 14.5 | 4.6 |
| AttnPacker | 0.96 | 0.64 | **1.15** | 31.52 | 32.98 | 53.16 | 55.48 | 50.1 | 83.4 | 22.5 | 7.2 |
| DLPacker | 1.01 | 0.65 | 1.22 | 32.59 | 32.39 | 55.95 | 72.09 | 50.1 | 77.3 | 12.9 | 3.0 |
| FASPR | 1.08 | 0.72 | 1.29 | 34.11 | 32.03 | 53.99 | 60.17 | 49.7 | 133.2 | 28.6 | 6.6 |
| PyRosetta Packer | 1.06 | 0.70 | 1.28 | 33.66 | 32.78 | 52.45 | 58.19 | 50.5 | 81.7 | 8.3 | 1.7 |
| SCWRL4 | 1.10 | 0.73 | 1.31 | 34.70 | 32.80 | 52.67 | 60.59 | 49.4 | 147.9 | 31.7 | 7.3 |

Values in bold indicate the best performance. Asterisks (\*) indicate values that are statistically significantly better than the corresponding value from AlphaFold baseline side-chain predictions at a 95% confidence level.

**Table 3.** Repacking performance of AlphaFold side-chains using a pLDDT-aware integrative approach for CASP15 targets (*n* = 71).

| | RMSD (Å) ↓ | | | χ-MAE (°) ↓ | | | | RR (%) ↑ | Steric Clashes (#) ↓ | | |
| --- | --- | --- | --- | --- | --- | --- | --- | --- | --- | --- | --- |
| | All | Core | Surface | $\chi_1$ | $\chi_2$ | $\chi_3$ | $\chi_4$ | $\chi_{1-4}$ | 100% | 90% | 80% |
| **AF2-Generated Backbones** |  |  |  |  |  |  |  |  |  |  |  |
| AlphaFold2 side-chains | 0.902 | 0.576 | 1.107 | 28.051 | 27.905 | **48.044** | 55.000 | 53.94 | 48.17 | 1.96 | **0.00** |
| Our integrative approach | **0.895** | **0.574** | **1.099** | **27.779** | **27.511** | 48.354 | **54.643** | **55.40** | **44.55** | **1.51** | 0.01 |
| *p*-values | <u>$3 \times 10^{-7}$</u> | 0.114 | <u>$1 \times 10^{-5}$</u> | <u>$7 \times 10^{-8}$</u> | <u>$2 \times 10^{-4}$</u> | 0.976 | <u>0.045</u> | <u>$7 \times 10^{-8}$</u> | <u>$3 \times 10^{-6}$</u> | <u>$4 \times 10^{-4}$</u> | 0.841 |
| **AF3-Generated Backbones** |  |  |  |  |  |  |  |  |  |  |  |
| AlphaFold3 side-chains | **0.951** | **0.600** | **1.159** | 30.182 | **28.902** | **48.920** | **53.937** | 53.83 | 58.39 | 8.07 | 1.01 |
| Our integrative approach | 0.954 | 0.605 | 1.159 | **30.177** | 29.066 | 49.698 | 54.540 | **54.07** | **50.73** | **4.30** | **0.24** |
| *p*-values | 0.913 | 0.896 | 0.541 | <u>0.047</u> | 0.687 | 0.997 | 0.996 | <u>0.036</u> | <u>$3 \times 10^{-11}$</u> | <u>$5 \times 10^{-11}$</u> | <u>0.002</u> |

Values in bold indicate the best performance. *p*-values are underlined if the method demonstrated statistically significantly better performance than AlphaFold for that metric at a 95% confidence level.

**Fig. 1.**
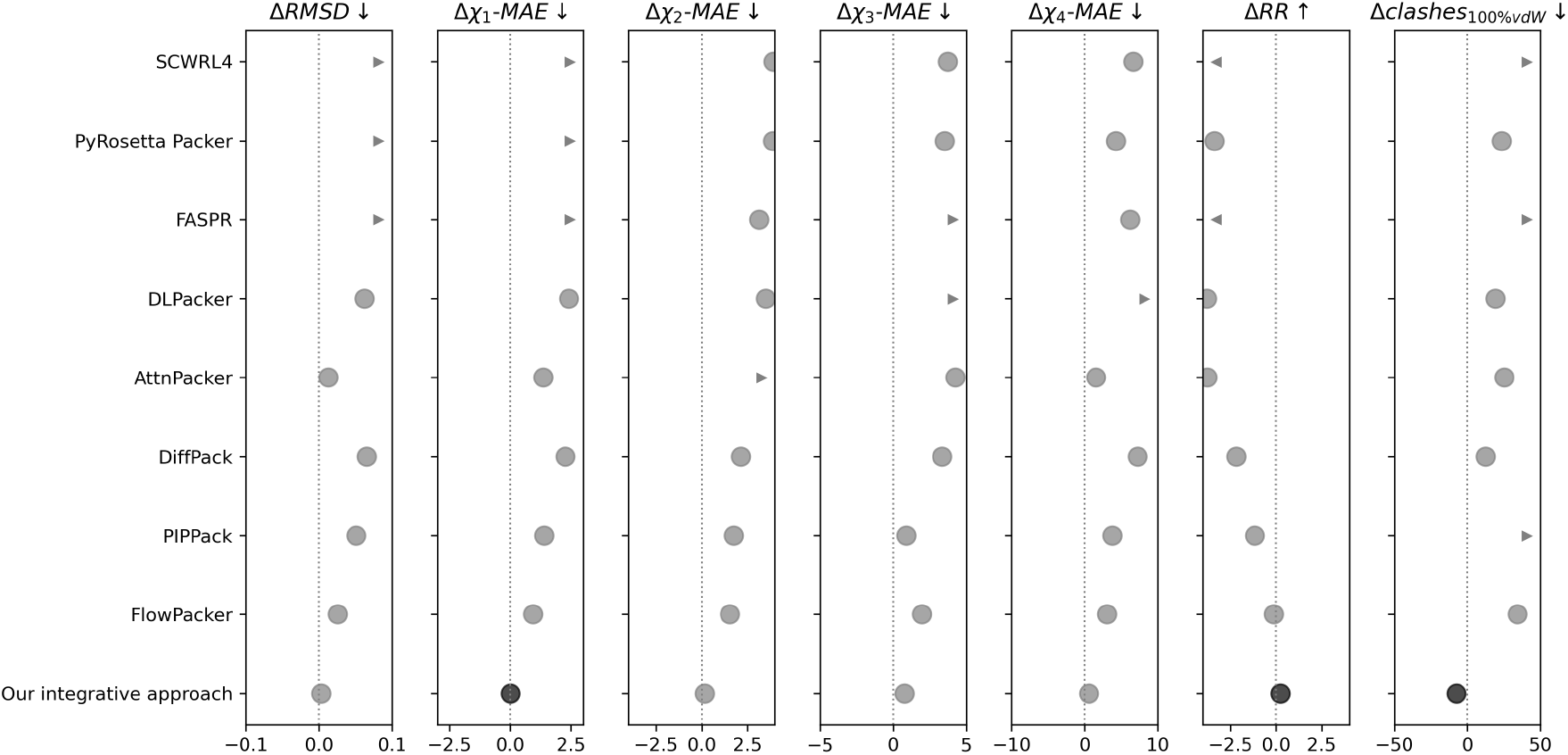
The performance difference between the baseline AlphaFold3-predicted side-chains and the repacked side-chains using various PSCP method for different evaluation metrics on the CASP15 targets. Filled black circles represent statistically significant improvements from AlphaFold3 at a 95% confidence level, whereas filled gray circles represent no significant improvement.

### 3.3. Repacking performance of AF side-chains using a backbone confidence-aware integrative approach

As shown in **Table 3**, our integrative approach that performs a weighted combination of several PSCP methods based on residue-level backbone pLDDT scores from AlphaFold attained modest yet statistically significant improvement (at a 95% confidence level) over the AlphaFold2 baseline performance for most evaluation metrics on CASP15 targets, having a rotameric recovery rate improvement of 1.5%. Compared to the AlphaFold3 baseline, however, there are negligible changes or even decreases in performance, with the exception of the recovery rate and mean absolute error in *χ*_1_ for which our integrative approach leads to minor but statistically significant improvement at a 95% confidence level. In summary, while our pLDDT-aware integrative approach often results in performance improvement over AlphaFold baselines, suggesting that plDDT can be useful for repacking side-chains when AlphaFold backbone coordinates are used as input instead of experimental input, it is still limited in its ability to yield consistent and significant improvement.

## 4. Conclusion

Here, we present a performance benchmarking of the current PSCP methods not just with experimental inputs, but also with AlphaFold2- and AlphaFold3-generated backbone coordinates as inputs. Empirical results demonstrate that while the PSCP methods are effective in packing the side-chains for experimental inputs, they fail to generalize in repacking AlphaFold-generated structures, consistently having side-chain accuracy lower than the AlphaFold baselines. We additionally explore the usefulness of leveraging the self-assessment confidence scores from AlphaFold in the form of residue-level backbone pLDDT by implementing a confidence-aware integrative approach that performs a weighted combination of several PSCP methods. The protocol often leads to performance gains by attaining modest yet statistically significant improvement, suggesting that plDDT can be useful for repacking side-chains AlphaFold backbone coordinates are used as input instead of experimental input. However, such a heuristic strategy is still far from consistent and significant improvement. Combining the power of the existing PSCP methods with residue-level backbone confidence estimation may be a promising avenue for further progress, thereby enhancing the fidelity of side-chain packing for improved protein structure prediction in the post-AlphaFold era.

## 5. Acknowledgments

This work was partially supported by the National Institute of General Medical Sciences (R35GM138146 to D.B.) and the National Science Foundation (DBI2208679 to D.B.).

