## Supplementary Information for "To pack or not to pack: revisiting protein side-chain packing in the post-AlphaFold era"

#### Contents

**1 Data Collection** **1**
1.1 Targets . . . . . . . . . . 1
1.2 Coordinate Data Generation . . . . . . . . . . 2

**2 Results from Assessment and Statistical Analysis** **3**

**3 Details of Our Algorithm for Repacking AlphaFold Side-Chains Using a Backbone Confidence-Aware Integrative Approach** **7**

### 1 Data Collection

#### 1.1 Targets

For our analyses, we curated a dataset for each of CASP14 and CASP15. The full lists of targets are given in **Table 1**. The following targets were removed from consideration:

- CASP14
  - The AlphaFold2 submissions in the CASP14 contests were missing the targets T1025, T1028, T1036s1, T1044, and T1045s1. As such, these were removed from our datasets.
- CASP15
  - T1169 was removed due to its length (3364 residues) causing high runtimes for our integrative energy optimization script.
  - T1186 was present in the list of native targets as well as AlphaFold2's publicly released predictions for CASP15, but it was not included in the list of sequences on the Protein Structure Prediction Center, so we could not generate AlphaFold3 predictions for it and therefore removed it from our study. Their website additionally labeled it as "Not a TS [tertiary structure] target: auxiliary structure for ligand prediction", though it was the

**Table 1.** List of targets used in this work.

| **Data Set** | **Target List** |
|---|---|
| CASP14 ($n = 66$) | T1024, T1026, T1027, T1029, T1030, T1031, T1032, T1033, T1034, T1035, T1037, T1038, T1039, T1040, T1041, T1042, T1043, T1045s2, T1046s1, T1046s2, T1047s1, T1047s2, T1048, T1049, T1050, T1052, T1053, T1054, T1055, T1056, T1057, T1058, T1060s2, T1060s3, T1061, T1062, T1064, T1065s1, T1065s2, T1067, T1068, T1070, T1072s1, T1073, T1074, T1076, T1078, T1079, T1080, T1082, T1083, T1084, T1087, T1088, T1089, T1090, T1091, T1092, T1093, T1094, T1095, T1096, T1098, T1099, T1100, T1101 |
| CASP15 ($n = 71$) | T1104, T1106s1, T1106s2, T1109, T1110, T1112, T1113, T1114s1, T1114s2, T1114s3, T1119, T1120, T1121, T1122, T1123, T1124, T1125, T1127, T1129s2, T1130, T1131, T1132, T1133, T1134s1, T1134s2, T1137s1, T1137s2, T1137s3, T1137s4, T1137s5, T1137s6, T1137s7, T1137s8, T1137s9, T1139, T1145, T1146, T1147, T1150, T1151s2, T1152, T1153, T1154, T1155, T1157s1, T1157s2, T1158, T1159, T1160, T1161, T1162, T1163, T1170, T1173, T1174, T1175, T1176, T1177, T1178, T1179, T1180, T1181, T1182, T1183, T1184, T1185s1, T1185s2, T1185s4, T1187, T1188, T1194 |

only target with a native structure that had this label, so no other targets were removed for this same reason.

#### 1.2 Coordinate Data Generation

From the sequences of the targets in CASP14 and CASP15, we used AlphaFold2 and AlphaFold3 to generate the predicted structures, the backbone coordinates of which are used in our study [1, 2]. All AlphaFold3 structures were generated using AlphaFold3's online server, located at `https://alphafoldserver.com/`, with a seed of 1048596 and converted from mmCIF to PDB with Project Gemmi's converter at `https://project-gemmi.github.io/wasm/convert/cif2pdb.html`. For both AF2 and AF3, 5 predictions were available for each target, from which we took the first for each.

For each side-chain packing method, we inputted the native structures, AF2-generated structures, and AF3-generated structures, and retrieved the tool's predictions. All tools were retrieved and installed approximately in September of 2024. Here are the relevant details for how we obtained the predicted structures from each side-chain packing tool:

**SCWRL4:** Inferences were made using the SCWRL4 [3] executable, downloaded from the Dunbrack Lab at `http://dunbrack.fccc.edu/lab/scwrl`.

**Rosetta Packer:** Our Rosetta Packer [4] predictions were obtained using PyRosetta's [5] `PackRotamersMover`, with a script largely derived from PIPPack's (`https://github.com/Kuhlman-Lab/PIPPack/blob/main/eval/rosetta_packer.py`).

**FASPR:** We compiled the FASPR [6] source code (found at `https://github.com/tommyhuangthu/FASPR`) into the executable and directly used that to generate each prediction.

One discrepancy to note is that FASPR did not initially run on targets T1041 and T1076 (only the native variants from CASP14) due to the very first atom in those PDBs (a nitrogen atom in the first serine amino acid residue) missing. So, we used the PULCHRA tool at `https://sites.gatech.edu/cssb/pulchra/` to restore any missing atoms and appended only that first nitrogen atom to the 2 respective PDB files. These mutated structures for the native CASP14 dataset were only used for FASPR to avoid involving unnecessary mutations for other tools.

**DLPacker:** As DLPacker [7] predicts one residue at a time and past predictions impact future ones for a given target, they developed 3 orderings of the residues in the structure to use, named `sequence`, `natoms`, and `score`. We ran all variants in our study.

**AttnPacker:** AttnPacker [8] was ran both with its default settings and with its post-processing procedure in a Jupyter Notebook.

**DiffPack:** DiffPack [9] was ran with its default configuration file and its configuration that used confidence sampling (`inference.yaml` and `inference_confidence.yaml` as found at `https://github.com/DeepGraphLearning/DiffPack/tree/main/config`, respectively). The random seed used was 1048596.

On the initial runs, DiffPack did not run on the following targets:

- *For native CASP14 backbones:* T1058, T1070, T1091, T1098
- *For AF2-predicted CASP14 backbones:* T1050, T1061, T1089
- *For native CASP15 backbones:* T1146, T1150, T1182
- *For AF3-predicted CASP15 backbones:* T1114s3, T1123, T1125, T1137s1, T1179

This list of non-conforming targets was shared by both the default and confidence sampling versions of DiffPack, and it was due to some missing atoms in the PDB files causing the explicit valence of specific atoms to be read as too large when parsing the PDBs. In order to alleviate this while still ensuring to use the original structures, the `sanitize` and `removeHs` parameters of the `rdkit.Chem.MolFromPDBFile()` function were both set to false (in order to allow DiffPack's `SideChainDataset.load_pdbs()` function to run), and the `strict` parameter of `mol.UpdatePropertyCache()` in TorchDrug's `Molecule.to_molecule()` method was also set to false.

**PIPPack:** The four variants of PIPPack [10] that were ran were: the default model, the default model with resampled coordinates, the ensembled inference model, and ensembled inferences with resampled coordinates.

**FlowPacker:** Four variations of FlowPacker [11] models were used: the `bc40.pth` checkpoint trained on BC40, that same checkpoint with confidence sampling (using `n_samples = 8`), the `cluster.pth` checkpoint trained on the PDB with a clustering threshold of 40% matching sequence identity, and that same checkpoint with confidence sampling (using `n_samples = 8`).

#### 2 Results from Assessment and Statistical Analysis

For evaluating all predicted structures, we compared them with the corresponding native structures using AttnPacker's protein side-chain assessment library, modifying it to categorize results among all residues, core residues, and surface residues, as well as to aggregate all targets' results together. The results are shown in **Tables 2** and **3**.

We ran a total of 792 Wilcoxon signed-rank tests for the original side-chain packing tools (2 CASPs $\times$ 2 AlphaFold versions to generate backbones $\times$ 18 total variations of tools $\times$ 11 metrics being evaluated). The results are given in **Tables 4** and **5**.

**Table 2.** Performance benchmarking on the CASP14 dataset (n = 66 targets).

| Input Structure | RMSD (Å) All | RMSD (Å) Core | RMSD (Å) Surface | $\chi$-MAE (°) $\chi_1$ | $\chi$-MAE (°) $\chi_2$ | $\chi$-MAE (°) $\chi_3$ | $\chi$-MAE (°) $\chi_4$ | $\chi$-Acc. (%) $\chi_{1-4}$ | Steric Clashes (#) 100% | Steric Clashes (#) 90% | Steric Clashes (#) 80% |
|---|---|---|---|---|---|---|---|---|---|---|---|
| **Sequence** | | | | | | | | | | | |
| AlphaFold2 | 1.07 | 0.66 | 1.26 | 34.59 | 31.51 | 50.81 | 51.42 | 46.0 | 41.4 | 1.9 | 0.0 |
| AlphaFold3 | 1.04 | 0.64 | 1.25 | 34.14 | 30.35 | 49.42 | 50.31 | 47.4 | 45.8 | 5.2 | 0.7 |
| **Native** | | | | | | | | | | | |
| FlowPacker$_{\mathrm{BC40}}$ | 0.83 | 0.43 | 1.04 | 23.80 | 26.17 | 47.06 | 53.56 | 56.2 | 116.7 | 27.8 | 8.3 |
| FlowPacker$_{\mathrm{BC40,\ conf}}$ | 0.83 | 0.42 | 1.05 | 23.62 | 26.28 | 46.82 | 55.01 | 56.6 | 118.2 | 28.3 | 8.7 |
| FlowPacker$_{\mathrm{cluster}}$ | 0.80 | 0.41 | 1.01 | 23.18 | 25.99 | 45.45 | 51.52 | 57.4 | 101.6 | 20.5 | 5.8 |
| FlowPacker$_{\mathrm{cluster,\ conf}}$ | 0.80 | 0.40 | 1.01 | 23.02 | 25.82 | 46.09 | 52.80 | 57.1 | 102.0 | 21.7 | 6.4 |
| PIPPack | 0.82 | 0.45 | 1.01 | 22.59 | 25.38 | 43.05 | 51.37 | 57.5 | 131.8 | 36.5 | 13.8 |
| PIPPack$_{\mathrm{RS}}$ | 0.82 | 0.45 | 1.01 | 22.65 | 25.59 | 43.50 | 51.05 | 57.3 | 111.1 | 23.8 | 6.7 |
| PIPPack$_{\mathrm{ensembled}}$ | 0.79 | 0.43 | 0.99 | 21.57 | 25.25 | 41.93 | 51.27 | 58.1 | 131.2 | 36.2 | 14.4 |
| PIPPack$_{\mathrm{ensembled\ +\ RS}}$ | 0.80 | 0.43 | 0.99 | 21.92 | 25.38 | 43.28 | 52.23 | 57.7 | 108.7 | 22.1 | 6.1 |
| DiffPack | 0.87 | 0.49 | 1.07 | 25.92 | 27.42 | 48.73 | 56.28 | 55.0 | 125.3 | 35.7 | 14.2 |
| DiffPack$_{\mathrm{conf}}$ | 0.79 | 0.41 | 0.98 | 22.92 | 25.23 | 46.97 | 55.33 | 57.6 | 104.2 | 26.8 | 9.8 |
| AttnPacker | 0.79 | 0.44 | 0.98 | 24.19 | 28.79 | 48.34 | 50.37 | 51.3 | 84.6 | 22.8 | 8.1 |
| AttnPacker$_{\mathrm{PP}}$ | 0.84 | 0.47 | 1.04 | 24.65 | 27.64 | 48.38 | 50.97 | 51.4 | 107.5 | 5.4 | 1.9 |
| DLPacker$_{\mathrm{seq}}$ | 0.92 | 0.52 | 1.13 | 27.93 | 30.31 | 52.35 | 70.46 | 49.6 | 93.9 | 20.9 | 7.7 |
| DLPacker$_{\mathrm{natoms}}$ | 0.92 | 0.52 | 1.12 | 27.98 | 30.38 | 54.25 | 73.34 | 49.5 | 88.0 | 18.2 | 5.8 |
| DLPacker$_{\mathrm{score}}$ | 0.90 | 0.50 | 1.11 | 27.45 | 30.03 | 52.82 | 70.34 | 50.6 | 83.2 | 16.8 | 5.1 |
| FASPR | 1.03 | 0.62 | 1.24 | 31.97 | 31.27 | 49.43 | 55.74 | 47.8 | 152.9 | 41.8 | 13.0 |
| PyRosetta | 1.00 | 0.55 | 1.23 | 30.98 | 31.29 | 49.31 | 55.58 | 48.9 | 104.3 | 22.1 | 8.4 |
| SCWRL4 | 1.04 | 0.61 | 1.26 | 32.22 | 31.65 | 50.21 | 55.10 | 47.5 | 158.3 | 40.2 | 11.8 |
| **AF2-Generated** | | | | | | | | | | | |
| FlowPacker$_{\mathrm{BC40}}$ | 1.09 | 0.67 | 1.30 | 35.51 | 33.13 | 53.66 | 59.77 | 45.5 | 84.8 | 13.2 | 2.6 |
| FlowPacker$_{\mathrm{BC40,\ conf}}$ | 1.09 | 0.67 | 1.31 | 35.65 | 33.18 | 53.25 | 57.32 | 45.5 | 85.2 | 12.8 | 2.5 |
| FlowPacker$_{\mathrm{cluster}}$ | 1.09 | 0.67 | 1.31 | 35.82 | 33.10 | 52.42 | 56.38 | 45.9 | 86.1 | 13.5 | 3.2 |
| FlowPacker$_{\mathrm{cluster,\ conf}}$ | 1.09 | 0.67 | 1.30 | 35.62 | 33.04 | 51.12 | 55.85 | 46.1 | 86.1 | 13.1 | 2.7 |
| PIPPack | 1.11 | 0.69 | 1.32 | 36.08 | 33.38 | 52.41 | 54.89 | 45.1 | 102.0 | 21.8 | 7.8 |
| PIPPack$_{\mathrm{RS}}$ | 1.11 | 0.69 | 1.33 | 36.17 | 33.59 | 53.13 | 54.73 | 44.7 | 91.0 | 14.6 | 3.1 |
| PIPPack$_{\mathrm{ensembled}}$ | 1.10 | 0.68 | 1.30 | 35.71 | 33.06 | 51.07 | 54.55 | 45.1 | 102.3 | 20.6 | 6.5 |
| PIPPack$_{\mathrm{ensembled\ +\ RS}}$ | 1.10 | 0.69 | 1.31 | 35.79 | 33.22 | 52.32 | 55.59 | 44.9 | 89.9 | 14.3 | 2.6 |
| DiffPack | 1.18 | 0.75 | 1.41 | 38.71 | 34.85 | 54.38 | 59.91 | 43.2 | 71.3 | 18.7 | 6.8 |
| DiffPack$_{\mathrm{conf}}$ | 1.12 | 0.70 | 1.34 | 36.69 | 32.99 | 53.41 | 56.13 | 44.9 | 57.2 | 11.7 | 3.7 |
| AttnPacker | 1.06 | 0.68 | 1.25 | 36.08 | 34.92 | 52.85 | 51.78 | 43.3 | 68.5 | 15.3 | 4.5 |
| AttnPacker$_{\mathrm{PP}}$ | 1.09 | 0.69 | 1.29 | 36.39 | 33.09 | 52.60 | 52.73 | 43.0 | 91.9 | 2.4 | 0.6 |
| DLPacker$_{\mathrm{seq}}$ | 1.11 | 0.70 | 1.32 | 36.44 | 35.41 | 57.26 | 73.90 | 42.8 | 75.1 | 13.3 | 2.7 |
| DLPacker$_{\mathrm{natoms}}$ | 1.12 | 0.70 | 1.33 | 36.82 | 35.40 | 57.80 | 73.11 | 42.5 | 71.3 | 12.2 | 2.9 |
| DLPacker$_{\mathrm{score}}$ | 1.11 | 0.70 | 1.33 | 36.28 | 35.19 | 57.87 | 72.25 | 42.9 | 67.7 | 11.0 | 2.1 |
| FASPR | 1.18 | 0.75 | 1.38 | 38.94 | 34.70 | 53.43 | 55.79 | 41.9 | 121.1 | 27.0 | 5.7 |
| PyRosetta | 1.16 | 0.72 | 1.38 | 38.16 | 35.19 | 52.84 | 55.31 | 42.7 | 73.9 | 7.7 | 1.2 |
| SCWRL4 | 1.20 | 0.77 | 1.40 | 39.01 | 35.09 | 52.93 | 56.20 | 41.6 | 132.8 | 29.0 | 5.7 |
| **AF3-Generated** | | | | | | | | | | | |
| FlowPacker$_{\mathrm{BC40}}$ | 1.08 | 0.67 | 1.29 | 35.22 | 32.14 | 51.64 | 54.21 | 46.4 | 81.1 | 13.0 | 2.8 |
| FlowPacker$_{\mathrm{BC40,\ conf}}$ | 1.08 | 0.67 | 1.29 | 34.91 | 32.35 | 51.33 | 54.67 | 46.9 | 79.3 | 11.8 | 2.4 |
| FlowPacker$_{\mathrm{cluster}}$ | 1.08 | 0.66 | 1.29 | 35.42 | 32.00 | 51.47 | 56.42 | 46.7 | 78.7 | 11.0 | 2.0 |
| FlowPacker$_{\mathrm{cluster,\ conf}}$ | 1.07 | 0.66 | 1.29 | 34.85 | 31.91 | 51.16 | 54.77 | 47.3 | 79.0 | 11.6 | 2.5 |
| PIPPack | 1.09 | 0.67 | 1.31 | 35.66 | 32.66 | 50.88 | 52.82 | 46.6 | 94.9 | 20.0 | 6.4 |
| PIPPack$_{\mathrm{RS}}$ | 1.09 | 0.67 | 1.31 | 35.77 | 32.78 | 51.17 | 53.40 | 46.3 | 82.7 | 12.8 | 2.2 |
| PIPPack$_{\mathrm{ensembled}}$ | 1.08 | 0.67 | 1.29 | 35.18 | 32.38 | 50.02 | 52.42 | 46.6 | 95.1 | 19.5 | 6.3 |
| PIPPack$_{\mathrm{ensembled\ +\ RS}}$ | 1.08 | 0.68 | 1.28 | 35.20 | 32.40 | 51.07 | 53.98 | 46.3 | 83.0 | 12.2 | 1.9 |
| DiffPack | 1.15 | 0.72 | 1.38 | 37.97 | 34.12 | 53.99 | 57.32 | 44.1 | 73.8 | 19.8 | 7.3 |
| DiffPack$_{\mathrm{conf}}$ | 1.11 | 0.69 | 1.33 | 36.43 | 32.66 | 51.65 | 56.08 | 45.8 | 56.0 | 12.1 | 4.0 |
| AttnPacker | 1.04 | 0.66 | 1.24* | 35.26 | 34.11 | 51.57 | 51.90 | 43.9 | 62.4 | 14.3 | 4.3 |
| AttnPacker$_{\mathrm{PP}}$ | 1.07 | 0.67 | 1.27 | 35.75 | 32.70 | 51.49 | 52.33 | 43.6 | 86.5 | 2.1* | 0.6 |
| DLPacker$_{\mathrm{seq}}$ | 1.11 | 0.69 | 1.33 | 36.42 | 34.95 | 56.12 | 70.69 | 43.6 | 68.2 | 11.4 | 3.1 |
| DLPacker$_{\mathrm{natoms}}$ | 1.10 | 0.70 | 1.31 | 36.17 | 34.60 | 56.69 | 73.09 | 43.2 | 65.3 | 9.8 | 2.0 |
| DLPacker$_{\mathrm{score}}$ | 1.10 | 0.69 | 1.31 | 36.28 | 34.13 | 56.91 | 72.26 | 43.3 | 64.5 | 10.3 | 2.2 |
| FASPR | 1.16 | 0.76 | 1.36 | 38.06 | 33.48 | 52.29 | 53.81 | 43.2 | 115.3 | 25.6 | 5.7 |
| PyRosetta | 1.15 | 0.71 | 1.37 | 37.66 | 34.51 | 52.23 | 54.16 | 43.4 | 69.3 | 7.1 | 1.4 |
| SCWRL4 | 1.17 | 0.77 | 1.37 | 37.69 | 34.12 | 51.83 | 57.18 | 43.1 | 127.7 | 28.1 | 6.4 |

Note: Asterisks (*) indicate values from repacking that are statistically significantly better than the corresponding value from AlphaFold's side-chain predictions.

**Table 3.** Performance benchmarking on the CASP15 dataset (n = 71 targets).

| | **RMSD (Å)** | | | **$\chi$-MAE (°)** | | | | **$\chi$-Acc. (%)** | **Steric Clashes (#)** | | |
|---|---|---|---|---|---|---|---|---|---|---|---|
| **Input Structure** | **All** | **Core** | **Surface** | $\chi_1$ | $\chi_2$ | $\chi_3$ | $\chi_4$ | $\chi_{1-4}$ | 100% | 90% | 80% |
| **Sequence** | | | | | | | | | | | |
| AlphaFold2 | 0.90 | 0.58 | 1.11 | 28.05 | 27.90 | 48.04 | 55.00 | 53.9 | 48.2 | 2.0 | 0.0 |
| AlphaFold3 | 0.95 | 0.60 | 1.16 | 30.18 | 28.90 | 48.92 | 53.94 | 53.8 | 58.4 | 8.1 | 1.0 |
| **Native** | | | | | | | | | | | |
| FlowPacker$_{\text{BC40}}$ | 0.73 | 0.35 | 0.95 | 20.27 | 23.43 | 43.18 | 54.89 | 64.7 | 111.5 | 18.4 | 3.9 |
| FlowPacker$_{\text{BC40, conf}}$ | 0.72 | 0.35 | 0.94 | 20.25 | 23.52 | 42.58 | 55.02 | 64.8 | 111.3 | 19.3 | 5.0 |
| FlowPacker$_{\text{cluster}}$ | 0.69 | 0.33 | 0.90 | 18.73 | 22.21 | 41.87 | 53.05 | 66.2 | 99.9 | 14.9 | 3.3 |
| FlowPacker$_{\text{cluster, conf}}$ | 0.69 | 0.33 | 0.90 | 18.99 | 22.04 | 40.93 | 52.62 | 66.4 | 100.8 | 14.6 | 3.3 |
| PIPPack | 0.72 | 0.35 | 0.93 | 19.05 | 22.70 | 41.13 | 51.95 | 65.7 | 128.5 | 30.7 | 11.5 |
| PIPPack$_{\text{RS}}$ | 0.73 | 0.35 | 0.94 | 19.23 | 22.81 | 41.65 | 52.14 | 65.2 | 108.1 | 18.3 | 3.8 |
| PIPPack$_{\text{ensembled}}$ | 0.70 | 0.34 | 0.91 | 18.27 | 22.16 | 40.21 | 53.36 | 66.1 | 129.0 | 30.5 | 10.9 |
| PIPPack$_{\text{ensembled + RS}}$ | 0.71 | 0.34 | 0.92 | 18.69 | 22.37 | 41.74 | 55.14 | 65.5 | 108.2 | 17.4 | 3.0 |
| DiffPack | 0.76 | 0.41 | 0.96 | 21.55 | 24.63 | 45.29 | 61.13 | 62.7 | 117.3 | 30.7 | 11.7 |
| DiffPack$_{\text{conf}}$ | 0.68 | 0.34 | 0.87 | 18.29 | 22.47 | 42.91 | 56.88 | 65.7 | 95.3 | 20.3 | 7.2 |
| AttnPacker | 0.71 | 0.37 | 0.90 | 20.29 | 26.10 | 47.09 | 54.68 | 59.2 | 96.4 | 25.5 | 9.5 |
| AttnPacker$_{\text{PP}}$ | 0.75 | 0.39 | 0.95 | 20.77 | 24.96 | 46.80 | 54.99 | 59.0 | 117.2 | 4.0 | 1.2 |
| DLPacker$_{\text{seq}}$ | 0.78 | 0.40 | 0.98 | 22.31 | 26.48 | 51.26 | 68.17 | 59.0 | 96.4 | 17.7 | 4.6 |
| DLPacker$_{\text{natoms}}$ | 0.77 | 0.40 | 0.98 | 22.39 | 26.08 | 50.99 | 68.79 | 59.2 | 92.3 | 15.4 | 3.6 |
| DLPacker$_{\text{score}}$ | 0.76 | 0.38 | 0.97 | 21.88 | 26.29 | 50.86 | 67.53 | 59.5 | 89.4 | 14.0 | 3.2 |
| FASPR | 0.92 | 0.52 | 1.14 | 27.12 | 29.07 | 50.39 | 59.05 | 55.8 | 160.5 | 37.4 | 9.7 |
| PyRosetta | 0.87 | 0.43 | 1.12 | 25.84 | 27.57 | 47.95 | 55.32 | 58.0 | 98.5 | 13.5 | 3.1 |
| SCWRL4 | 0.94 | 0.50 | 1.17 | 27.89 | 29.12 | 49.81 | 57.25 | 55.5 | 168.3 | 36.3 | 7.7 |
| **AF2-Generated** | | | | | | | | | | | |
| FlowPacker$_{\text{BC40}}$ | 0.95 | 0.59 | 1.16 | 30.10 | 29.87 | 50.56 | 60.66 | 53.5 | 104.2 | 15.7 | 3.0 |
| FlowPacker$_{\text{BC40, conf}}$ | 0.95 | 0.60 | 1.17 | 29.84 | 29.66 | 50.64 | 61.77 | 53.7 | 104.2 | 15.5 | 3.0 |
| FlowPacker$_{\text{cluster}}$ | 0.94 | 0.59 | 1.16 | 29.80 | 29.33 | 49.20 | 56.43 | 54.9* | 102.3 | 14.4 | 2.8 |
| FlowPacker$_{\text{cluster, conf}}$ | 0.94 | 0.59 | 1.15 | 29.38 | 29.18 | 50.18 | 57.00 | 55.1* | 100.4 | 13.9 | 2.3 |
| PIPPack | 0.97 | 0.60 | 1.20 | 30.55 | 30.00 | 51.48 | 57.15 | 53.4 | 123.9 | 25.6 | 8.2 |
| PIPPack$_{\text{RS}}$ | 0.98 | 0.62 | 1.20 | 30.66 | 30.53 | 52.80 | 57.51 | 53.1 | 109.3 | 17.2 | 3.5 |
| PIPPack$_{\text{ensembled}}$ | 0.96 | 0.61 | 1.19 | 30.08 | 29.89 | 50.22 | 56.32 | 53.5 | 124.5 | 27.3 | 9.4 |
| PIPPack$_{\text{ensembled + RS}}$ | 0.97 | 0.62 | 1.19 | 30.31 | 30.25 | 51.20 | 58.68 | 52.8 | 107.7 | 16.8 | 3.1 |
| DiffPack | 1.04 | 0.65 | 1.28 | 32.75 | 31.82 | 53.60 | 58.29 | 51.0 | 91.6 | 24.4 | 9.5 |
| DiffPack$_{\text{conf}}$ | 0.99 | 0.63 | 1.21 | 31.07 | 30.10 | 51.90 | 56.39 | 52.8 | 69.7 | 13.7 | 4.5 |
| AttnPacker | 0.92 | 0.61 | 1.12 | 30.34 | 31.48 | 52.10 | 55.52 | 50.4 | 87.9 | 23.6 | 7.7 |
| AttnPacker$_{\text{PP}}$ | 0.96 | 0.64 | 1.17 | 30.62 | 30.04 | 51.97 | 56.22 | 50.4 | 112.2 | 3.2 | 0.8 |
| DLPacker$_{\text{seq}}$ | 0.99 | 0.63 | 1.20 | 31.55 | 32.20 | 56.79 | 71.11 | 50.4 | 89.3 | 15.3 | 3.5 |
| DLPacker$_{\text{natoms}}$ | 0.97 | 0.64 | 1.18 | 31.08 | 31.48 | 56.02 | 71.83 | 50.6 | 85.5 | 13.7 | 2.8 |
| DLPacker$_{\text{score}}$ | 0.97 | 0.62 | 1.18 | 30.94 | 31.05 | 55.90 | 69.34 | 51.3 | 84.2 | 12.9 | 2.7 |
| FASPR | 1.06 | 0.71 | 1.27 | 33.67 | 32.53 | 53.56 | 59.75 | 50.0 | 147.0 | 32.5 | 7.9 |
| PyRosetta | 1.04 | 0.67 | 1.26 | 32.78 | 32.67 | 51.42 | 58.65 | 51.0 | 91.6 | 10.7 | 2.4 |
| SCWRL4 | 1.07 | 0.71 | 1.29 | 33.95 | 32.73 | 53.35 | 61.55 | 50.0 | 160.7 | 35.0 | 7.5 |
| **AF3-Generated** | | | | | | | | | | | |
| FlowPacker$_{\text{BC40}}$ | 1.00 | 0.62 | 1.22 | 31.81 | 31.12 | 50.02 | 59.79 | 52.9 | 96.1 | 16.1 | 3.8 |
| FlowPacker$_{\text{BC40, conf}}$ | 0.99 | 0.64 | 1.21 | 31.48 | 30.79 | 50.54 | 59.04 | 53.0 | 97.2 | 15.5 | 3.7 |
| FlowPacker$_{\text{cluster}}$ | 0.98 | 0.61 | 1.20 | 31.55 | 30.27 | 49.23 | 56.57 | 54.0 | 91.5 | 13.5 | 2.4 |
| FlowPacker$_{\text{cluster, conf}}$ | 0.98 | 0.62 | 1.19 | 31.12 | 30.44 | 50.88 | 57.02 | 53.7 | 92.5 | 13.3 | 2.5 |
| PIPPack | 1.01 | 0.65 | 1.23 | 31.84 | 31.24 | 50.23 | 58.05 | 52.8 | 115.1 | 24.9 | 8.4 |
| PIPPack$_{\text{RS}}$ | 1.02 | 0.64 | 1.24 | 31.95 | 31.38 | 51.13 | 58.23 | 52.6 | 98.9 | 16.0 | 3.0 |
| PIPPack$_{\text{ensembled}}$ | 1.00 | 0.64 | 1.22 | 31.57 | 30.65 | 49.81 | 57.75 | 52.7 | 115.7 | 26.5 | 9.0 |
| PIPPack$_{\text{ensembled + RS}}$ | 1.00 | 0.65 | 1.22 | 31.65 | 30.78 | 51.20 | 58.12 | 52.1 | 98.2 | 14.9 | 2.7 |
| DiffPack | 1.07 | 0.70 | 1.30 | 33.76 | 33.22 | 54.93 | 62.44 | 50.1 | 85.7 | 22.0 | 7.9 |
| DiffPack$_{\text{conf}}$ | 1.02 | 0.66 | 1.23 | 32.44 | 31.03 | 52.27 | 61.20 | 51.6 | 70.8 | 14.5 | 4.6 |
| AttnPacker | 0.96 | 0.64 | 1.15 | 31.52 | 32.98 | 53.16 | 55.48 | 50.1 | 83.4 | 22.5 | 7.2 |
| AttnPacker$_{\text{PP}}$ | 1.00 | 0.66 | 1.20 | 31.90 | 31.34 | 53.14 | 55.79 | 50.3 | 104.0 | 2.5* | 0.7 |
| DLPacker$_{\text{seq}}$ | 1.02 | 0.67 | 1.23 | 32.65 | 33.52 | 56.95 | 72.58 | 49.4 | 81.8 | 15.0 | 3.9 |
| DLPacker$_{\text{natoms}}$ | 1.03 | 0.68 | 1.23 | 33.12 | 33.04 | 56.17 | 71.49 | 49.6 | 79.3 | 12.5 | 2.5 |
| DLPacker$_{\text{score}}$ | 1.01 | 0.65 | 1.22 | 32.59 | 32.39 | 55.95 | 72.09 | 50.1 | 77.3 | 12.9 | 3.0 |
| FASPR | 1.08 | 0.72 | 1.29 | 34.11 | 32.03 | 53.99 | 60.17 | 49.7 | 133.2 | 28.6 | 6.6 |
| PyRosetta | 1.06 | 0.70 | 1.28 | 33.66 | 32.78 | 52.45 | 58.19 | 50.5 | 81.7 | 8.3 | 1.7 |
| SCWRL4 | 1.10 | 0.73 | 1.31 | 34.70 | 32.80 | 52.67 | 60.59 | 49.4 | 147.9 | 31.7 | 7.3 |

Note: Asterisks (*) indicate values from repacking that are statistically significantly better than the corresponding value from AlphaFold's side-chain predictions.

**Table 4.** Statistical significance test results ($p$-values) for the CASP14 dataset (n = 66 targets).

| | **RMSD (Å)** | | | **$\chi$-MAE (°)** | | | | **$\chi$-Acc. (%)** | **Steric Clashes (#)** | | |
|---|---|---|---|---|---|---|---|---|---|---|---|
| **Repacking Tool** | **All** | **Core** | **Surface** | $\chi_1$ | $\chi_2$ | $\chi_3$ | $\chi_4$ | $\chi_{1-4}$ | 100% | 90% | 80% |
| **Repacking AlphaFold2's Predicted Backbones** | | | | | | | | | | | |
| FlowPacker$_{BC40}$ | $9.9992 \times 10^{-1}$ | $6.2674 \times 10^{-1}$ | $9.9980 \times 10^{-1}$ | $9.9957 \times 10^{-1}$ | $9.9999 \times 10^{-1}$ | $9.9883 \times 10^{-1}$ | $9.9999 \times 10^{-1}$ | $7.8328 \times 10^{-1}$ | $9.9999 \times 10^{-1}$ | $9.9999 \times 10^{-1}$ | $9.9999 \times 10^{-1}$ |
| FlowPacker$_{BC40, conf}$ | $9.9996 \times 10^{-1}$ | $7.6072 \times 10^{-1}$ | $9.9993 \times 10^{-1}$ | $9.9940 \times 10^{-1}$ | $9.9997 \times 10^{-1}$ | $9.9878 \times 10^{-1}$ | $9.9985 \times 10^{-1}$ | $8.0310 \times 10^{-1}$ | $9.9999 \times 10^{-1}$ | $9.9999 \times 10^{-1}$ | $9.9999 \times 10^{-1}$ |
| FlowPacker$_{cluster}$ | $9.9979 \times 10^{-1}$ | $5.0839 \times 10^{-1}$ | $9.9988 \times 10^{-1}$ | $9.9989 \times 10^{-1}$ | $9.9999 \times 10^{-1}$ | $9.6959 \times 10^{-1}$ | $9.9989 \times 10^{-1}$ | $7.9751 \times 10^{-1}$ | $9.9999 \times 10^{-1}$ | $9.9999 \times 10^{-1}$ | $9.9999 \times 10^{-1}$ |
| FlowPacker$_{cluster, conf}$ | $9.9939 \times 10^{-1}$ | $6.5626 \times 10^{-1}$ | $9.9990 \times 10^{-1}$ | $9.9935 \times 10^{-1}$ | $9.9997 \times 10^{-1}$ | $8.8324 \times 10^{-1}$ | $9.9890 \times 10^{-1}$ | $4.0814 \times 10^{-1}$ | $9.9999 \times 10^{-1}$ | $9.9999 \times 10^{-1}$ | $9.9999 \times 10^{-1}$ |
| PIPPack | $9.9999 \times 10^{-1}$ | $9.9767 \times 10^{-1}$ | $9.9999 \times 10^{-1}$ | $9.9999 \times 10^{-1}$ | $9.9999 \times 10^{-1}$ | $9.7669 \times 10^{-1}$ | $9.9449 \times 10^{-1}$ | $9.5867 \times 10^{-1}$ | $9.9999 \times 10^{-1}$ | $9.9999 \times 10^{-1}$ | $9.9999 \times 10^{-1}$ |
| PIPPack$_{RS}$ | $9.9999 \times 10^{-1}$ | $9.9918 \times 10^{-1}$ | $9.9999 \times 10^{-1}$ | $9.9999 \times 10^{-1}$ | $9.9999 \times 10^{-1}$ | $9.9690 \times 10^{-1}$ | $9.9880 \times 10^{-1}$ | $9.9637 \times 10^{-1}$ | $9.9999 \times 10^{-1}$ | $9.9999 \times 10^{-1}$ | $9.9999 \times 10^{-1}$ |
| PIPPack$_{ensembled}$ | $9.9999 \times 10^{-1}$ | $9.9549 \times 10^{-1}$ | $9.9995 \times 10^{-1}$ | $9.9988 \times 10^{-1}$ | $9.9998 \times 10^{-1}$ | $8.0134 \times 10^{-1}$ | $9.6238 \times 10^{-1}$ | $9.3851 \times 10^{-1}$ | $9.9999 \times 10^{-1}$ | $9.9999 \times 10^{-1}$ | $9.9999 \times 10^{-1}$ |
| PIPPack$_{ensembled + RS}$ | $9.9999 \times 10^{-1}$ | $9.9988 \times 10^{-1}$ | $9.9996 \times 10^{-1}$ | $9.9994 \times 10^{-1}$ | $9.9999 \times 10^{-1}$ | $9.8494 \times 10^{-1}$ | $9.9924 \times 10^{-1}$ | $9.8397 \times 10^{-1}$ | $9.9999 \times 10^{-1}$ | $9.9999 \times 10^{-1}$ | $9.9999 \times 10^{-1}$ |
| DiffPack | $9.9999 \times 10^{-1}$ | $9.9999 \times 10^{-1}$ | $9.9999 \times 10^{-1}$ | $9.9999 \times 10^{-1}$ | $9.9999 \times 10^{-1}$ | $9.9994 \times 10^{-1}$ | $9.9999 \times 10^{-1}$ | $9.9999 \times 10^{-1}$ | $9.9999 \times 10^{-1}$ | $9.9999 \times 10^{-1}$ | $9.9999 \times 10^{-1}$ |
| DiffPack$_{conf}$ | $9.9999 \times 10^{-1}$ | $9.9998 \times 10^{-1}$ | $9.9999 \times 10^{-1}$ | $9.9999 \times 10^{-1}$ | $9.9998 \times 10^{-1}$ | $9.9976 \times 10^{-1}$ | $9.9989 \times 10^{-1}$ | $9.9733 \times 10^{-1}$ | $9.9999 \times 10^{-1}$ | $9.9999 \times 10^{-1}$ | $9.9999 \times 10^{-1}$ |
| AttnPacker | $3.7087 \times 10^{-1}$ | $9.0392 \times 10^{-1}$ | $5.0980 \times 10^{-2}$ | $9.9999 \times 10^{-1}$ | $9.9999 \times 10^{-1}$ | $9.9872 \times 10^{-1}$ | $6.6683 \times 10^{-1}$ | $9.9999 \times 10^{-1}$ | $9.9999 \times 10^{-1}$ | $9.9999 \times 10^{-1}$ | $9.9999 \times 10^{-1}$ |
| AttnPacker$_{PP}$ | $9.9999 \times 10^{-1}$ | $9.9850 \times 10^{-1}$ | $9.9933 \times 10^{-1}$ | $9.9999 \times 10^{-1}$ | $9.9999 \times 10^{-1}$ | $9.9867 \times 10^{-1}$ | $8.7291 \times 10^{-1}$ | $9.9999 \times 10^{-1}$ | $9.9999 \times 10^{-1}$ | $4.8302 \times 10^{-1}$ | $9.9971 \times 10^{-1}$ |
| DLPacker$_{seq}$ | $9.9999 \times 10^{-1}$ | $9.9998 \times 10^{-1}$ | $9.9999 \times 10^{-1}$ | $9.9999 \times 10^{-1}$ | $9.9999 \times 10^{-1}$ | $9.9999 \times 10^{-1}$ | $9.9999 \times 10^{-1}$ | $9.9999 \times 10^{-1}$ | $9.9999 \times 10^{-1}$ | $9.9999 \times 10^{-1}$ | $9.9999 \times 10^{-1}$ |
| DLPacker$_{natoms}$ | $9.9999 \times 10^{-1}$ | $9.9997 \times 10^{-1}$ | $9.9999 \times 10^{-1}$ | $9.9999 \times 10^{-1}$ | $9.9999 \times 10^{-1}$ | $9.9999 \times 10^{-1}$ | $9.9999 \times 10^{-1}$ | $9.9999 \times 10^{-1}$ | $9.9999 \times 10^{-1}$ | $9.9999 \times 10^{-1}$ | $9.9999 \times 10^{-1}$ |
| DLPacker$_{score}$ | $9.9999 \times 10^{-1}$ | $9.9996 \times 10^{-1}$ | $9.9999 \times 10^{-1}$ | $9.9999 \times 10^{-1}$ | $9.9999 \times 10^{-1}$ | $9.9999 \times 10^{-1}$ | $9.9999 \times 10^{-1}$ | $9.9999 \times 10^{-1}$ | $9.9999 \times 10^{-1}$ | $9.9999 \times 10^{-1}$ | $9.9999 \times 10^{-1}$ |
| FASPR | $9.9999 \times 10^{-1}$ | $9.9999 \times 10^{-1}$ | $9.9999 \times 10^{-1}$ | $9.9999 \times 10^{-1}$ | $9.9999 \times 10^{-1}$ | $9.9989 \times 10^{-1}$ | $9.9978 \times 10^{-1}$ | $9.9999 \times 10^{-1}$ | $9.9999 \times 10^{-1}$ | $9.9999 \times 10^{-1}$ | $9.9999 \times 10^{-1}$ |
| PyRosetta | $9.9999 \times 10^{-1}$ | $9.9996 \times 10^{-1}$ | $9.9999 \times 10^{-1}$ | $9.9999 \times 10^{-1}$ | $9.9999 \times 10^{-1}$ | $9.9872 \times 10^{-1}$ | $9.9905 \times 10^{-1}$ | $9.9999 \times 10^{-1}$ | $9.9999 \times 10^{-1}$ | $9.9999 \times 10^{-1}$ | $9.9999 \times 10^{-1}$ |
| SCWRL4 | $9.9999 \times 10^{-1}$ | $9.9999 \times 10^{-1}$ | $9.9999 \times 10^{-1}$ | $9.9999 \times 10^{-1}$ | $9.9999 \times 10^{-1}$ | $9.9791 \times 10^{-1}$ | $9.9976 \times 10^{-1}$ | $9.9999 \times 10^{-1}$ | $9.9999 \times 10^{-1}$ | $9.9999 \times 10^{-1}$ | $9.9999 \times 10^{-1}$ |
| **Repacking AlphaFold3's Predicted Backbones** | | | | | | | | | | | |
| FlowPacker$_{BC40}$ | $9.9999 \times 10^{-1}$ | $9.9997 \times 10^{-1}$ | $9.9946 \times 10^{-1}$ | $9.9972 \times 10^{-1}$ | $9.9999 \times 10^{-1}$ | $9.9954 \times 10^{-1}$ | $9.9943 \times 10^{-1}$ | $9.9584 \times 10^{-1}$ | $9.9999 \times 10^{-1}$ | $9.9999 \times 10^{-1}$ | $9.9999 \times 10^{-1}$ |
| FlowPacker$_{BC40, conf}$ | $9.9999 \times 10^{-1}$ | $9.9996 \times 10^{-1}$ | $9.9979 \times 10^{-1}$ | $9.9917 \times 10^{-1}$ | $9.9999 \times 10^{-1}$ | $9.9927 \times 10^{-1}$ | $9.9974 \times 10^{-1}$ | $9.7427 \times 10^{-1}$ | $9.9999 \times 10^{-1}$ | $9.9999 \times 10^{-1}$ | $9.9999 \times 10^{-1}$ |
| FlowPacker$_{cluster}$ | $9.9999 \times 10^{-1}$ | $9.9806 \times 10^{-1}$ | $9.9998 \times 10^{-1}$ | $9.9997 \times 10^{-1}$ | $9.9999 \times 10^{-1}$ | $9.9979 \times 10^{-1}$ | $9.9999 \times 10^{-1}$ | $9.9366 \times 10^{-1}$ | $9.9999 \times 10^{-1}$ | $9.9999 \times 10^{-1}$ | $9.9993 \times 10^{-1}$ |
| FlowPacker$_{cluster, conf}$ | $9.9999 \times 10^{-1}$ | $9.8589 \times 10^{-1}$ | $9.9991 \times 10^{-1}$ | $9.9511 \times 10^{-1}$ | $9.9996 \times 10^{-1}$ | $9.9538 \times 10^{-1}$ | $9.9966 \times 10^{-1}$ | $9.2206 \times 10^{-1}$ | $9.9999 \times 10^{-1}$ | $9.9999 \times 10^{-1}$ | $9.9999 \times 10^{-1}$ |
| PIPPack | $9.9999 \times 10^{-1}$ | $9.9997 \times 10^{-1}$ | $9.9999 \times 10^{-1}$ | $9.9996 \times 10^{-1}$ | $9.9999 \times 10^{-1}$ | $9.9963 \times 10^{-1}$ | $9.7046 \times 10^{-1}$ | $9.9840 \times 10^{-1}$ | $9.9999 \times 10^{-1}$ | $9.9999 \times 10^{-1}$ | $9.9999 \times 10^{-1}$ |
| PIPPack$_{RS}$ | $9.9999 \times 10^{-1}$ | $9.9999 \times 10^{-1}$ | $9.9999 \times 10^{-1}$ | $9.9999 \times 10^{-1}$ | $9.9999 \times 10^{-1}$ | $9.9943 \times 10^{-1}$ | $9.9071 \times 10^{-1}$ | $9.9970 \times 10^{-1}$ | $9.9999 \times 10^{-1}$ | $9.9999 \times 10^{-1}$ | $9.9996 \times 10^{-1}$ |
| PIPPack$_{ensembled}$ | $9.9999 \times 10^{-1}$ | $9.9999 \times 10^{-1}$ | $9.9960 \times 10^{-1}$ | $9.9907 \times 10^{-1}$ | $9.9999 \times 10^{-1}$ | $9.9297 \times 10^{-1}$ | $9.8542 \times 10^{-1}$ | $9.9520 \times 10^{-1}$ | $9.9999 \times 10^{-1}$ | $9.9999 \times 10^{-1}$ | $9.9999 \times 10^{-1}$ |
| PIPPack$_{ensembled + RS}$ | $9.9999 \times 10^{-1}$ | $9.9999 \times 10^{-1}$ | $9.9929 \times 10^{-1}$ | $9.9965 \times 10^{-1}$ | $9.9999 \times 10^{-1}$ | $9.9903 \times 10^{-1}$ | $9.9929 \times 10^{-1}$ | $9.9952 \times 10^{-1}$ | $9.9999 \times 10^{-1}$ | $9.9999 \times 10^{-1}$ | $9.9907 \times 10^{-1}$ |
| DiffPack | $9.9999 \times 10^{-1}$ | $9.9999 \times 10^{-1}$ | $9.9999 \times 10^{-1}$ | $9.9999 \times 10^{-1}$ | $9.9999 \times 10^{-1}$ | $9.9999 \times 10^{-1}$ | $9.9999 \times 10^{-1}$ | $9.9999 \times 10^{-1}$ | $9.9999 \times 10^{-1}$ | $9.9999 \times 10^{-1}$ | $9.9999 \times 10^{-1}$ |
| DiffPack$_{conf}$ | $9.9999 \times 10^{-1}$ | $9.9999 \times 10^{-1}$ | $9.9999 \times 10^{-1}$ | $9.9999 \times 10^{-1}$ | $9.9999 \times 10^{-1}$ | $9.9845 \times 10^{-1}$ | $9.9999 \times 10^{-1}$ | $9.9999 \times 10^{-1}$ | $9.9994 \times 10^{-1}$ | $9.9999 \times 10^{-1}$ | $9.9999 \times 10^{-1}$ |
| AttnPacker | $5.8473 \times 10^{-1}$ | $9.9933 \times 10^{-1}$ | $2.7445 \times 10^{-2}$* | $9.9901 \times 10^{-1}$ | $9.9999 \times 10^{-1}$ | $9.9870 \times 10^{-1}$ | $9.7003 \times 10^{-1}$ | $9.9999 \times 10^{-1}$ | $9.9999 \times 10^{-1}$ | $9.9999 \times 10^{-1}$ | $9.9999 \times 10^{-1}$ |
| AttnPacker$_{PP}$ | $9.9999 \times 10^{-1}$ | $9.9998 \times 10^{-1}$ | $9.8871 \times 10^{-1}$ | $9.9999 \times 10^{-1}$ | $9.9999 \times 10^{-1}$ | $9.9791 \times 10^{-1}$ | $9.8880 \times 10^{-1}$ | $9.9999 \times 10^{-1}$ | $9.9999 \times 10^{-1}$ | $2.6459 \times 10^{-7}$* | $4.4392 \times 10^{-1}$ |
| DLPacker$_{seq}$ | $9.9999 \times 10^{-1}$ | $9.9999 \times 10^{-1}$ | $9.9999 \times 10^{-1}$ | $9.9999 \times 10^{-1}$ | $9.9999 \times 10^{-1}$ | $9.9999 \times 10^{-1}$ | $9.9999 \times 10^{-1}$ | $9.9999 \times 10^{-1}$ | $9.9999 \times 10^{-1}$ | $9.9999 \times 10^{-1}$ | $9.9999 \times 10^{-1}$ |
| DLPacker$_{natoms}$ | $9.9999 \times 10^{-1}$ | $9.9999 \times 10^{-1}$ | $9.9999 \times 10^{-1}$ | $9.9999 \times 10^{-1}$ | $9.9999 \times 10^{-1}$ | $9.9999 \times 10^{-1}$ | $9.9999 \times 10^{-1}$ | $9.9999 \times 10^{-1}$ | $9.9999 \times 10^{-1}$ | $9.9999 \times 10^{-1}$ | $9.9999 \times 10^{-1}$ |
| DLPacker$_{score}$ | $9.9999 \times 10^{-1}$ | $9.9999 \times 10^{-1}$ | $9.9999 \times 10^{-1}$ | $9.9999 \times 10^{-1}$ | $9.9999 \times 10^{-1}$ | $9.9999 \times 10^{-1}$ | $9.9999 \times 10^{-1}$ | $9.9999 \times 10^{-1}$ | $9.9999 \times 10^{-1}$ | $9.9999 \times 10^{-1}$ | $9.9999 \times 10^{-1}$ |
| FASPR | $9.9999 \times 10^{-1}$ | $9.9999 \times 10^{-1}$ | $9.9999 \times 10^{-1}$ | $9.9999 \times 10^{-1}$ | $9.9999 \times 10^{-1}$ | $9.9933 \times 10^{-1}$ | $9.9897 \times 10^{-1}$ | $9.9999 \times 10^{-1}$ | $9.9999 \times 10^{-1}$ | $9.9999 \times 10^{-1}$ | $9.9999 \times 10^{-1}$ |
| PyRosetta | $9.9999 \times 10^{-1}$ | $9.9999 \times 10^{-1}$ | $9.9999 \times 10^{-1}$ | $9.9999 \times 10^{-1}$ | $9.9999 \times 10^{-1}$ | $9.9950 \times 10^{-1}$ | $9.9978 \times 10^{-1}$ | $9.9999 \times 10^{-1}$ | $9.9999 \times 10^{-1}$ | $9.9381 \times 10^{-1}$ | $9.8180 \times 10^{-1}$ |
| SCWRL4 | $9.9999 \times 10^{-1}$ | $9.9999 \times 10^{-1}$ | $9.9999 \times 10^{-1}$ | $9.9999 \times 10^{-1}$ | $9.9999 \times 10^{-1}$ | $9.9951 \times 10^{-1}$ | $9.9999 \times 10^{-1}$ | $9.9999 \times 10^{-1}$ | $9.9999 \times 10^{-1}$ | $9.9999 \times 10^{-1}$ | $9.9999 \times 10^{-1}$ |

Note: Asterisks (*) indicate values that are statistically significantly better than the corresponding value from AlphaFold's side-chain predictions.

**Table 5.** Statistical significance test results ($p$-values) for the CASP15 dataset (n = 71 targets).

| | **RMSD (Å)** | | | **$\chi$-MAE (°)** | | | | **$\chi$-Acc. (%)** | **Steric Clashes (#)** | | |
|---|---|---|---|---|---|---|---|---|---|---|---|
| **Repacking Tool** | **All** | **Core** | **Surface** | $\chi_1$ | $\chi_2$ | $\chi_3$ | $\chi_4$ | $\chi_{1-4}$ | 100% | 90% | 80% |
| **Repacking AlphaFold2's Predicted Backbones** | | | | | | | | | | | |
| FlowPacker$_{BC40}$ | $9.9999 \times 10^{-1}$ | $9.7824 \times 10^{-1}$ | $9.9999 \times 10^{-1}$ | $9.9999 \times 10^{-1}$ | $9.9999 \times 10^{-1}$ | $9.9968 \times 10^{-1}$ | $9.9997 \times 10^{-1}$ | $7.4883 \times 10^{-1}$ | $9.9999 \times 10^{-1}$ | $9.9999 \times 10^{-1}$ | $9.9999 \times 10^{-1}$ |
| FlowPacker$_{BC40, conf}$ | $9.9999 \times 10^{-1}$ | $9.3187 \times 10^{-1}$ | $9.9999 \times 10^{-1}$ | $9.9999 \times 10^{-1}$ | $9.9999 \times 10^{-1}$ | $9.9982 \times 10^{-1}$ | $9.9999 \times 10^{-1}$ | $6.7066 \times 10^{-1}$ | $9.9999 \times 10^{-1}$ | $9.9999 \times 10^{-1}$ | $9.9999 \times 10^{-1}$ |
| FlowPacker$_{cluster}$ | $9.9998 \times 10^{-1}$ | $9.3060 \times 10^{-1}$ | $9.9997 \times 10^{-1}$ | $9.9999 \times 10^{-1}$ | $9.9998 \times 10^{-1}$ | $9.5666 \times 10^{-1}$ | $8.0178 \times 10^{-1}$ | $1.9579 \times 10^{-2}$* | $9.9999 \times 10^{-1}$ | $9.9999 \times 10^{-1}$ | $9.9999 \times 10^{-1}$ |
| FlowPacker$_{cluster, conf}$ | $9.9993 \times 10^{-1}$ | $5.9715 \times 10^{-1}$ | $9.9992 \times 10^{-1}$ | $9.9985 \times 10^{-1}$ | $9.9998 \times 10^{-1}$ | $9.9607 \times 10^{-1}$ | $9.5692 \times 10^{-1}$ | $6.0956 \times 10^{-3}$* | $9.9999 \times 10^{-1}$ | $9.9999 \times 10^{-1}$ | $9.9999 \times 10^{-1}$ |
| PIPPack | $9.9999 \times 10^{-1}$ | $9.9988 \times 10^{-1}$ | $9.9999 \times 10^{-1}$ | $9.9999 \times 10^{-1}$ | $9.9999 \times 10^{-1}$ | $9.9997 \times 10^{-1}$ | $9.6789 \times 10^{-1}$ | $9.5387 \times 10^{-1}$ | $9.9999 \times 10^{-1}$ | $9.9999 \times 10^{-1}$ | $9.9999 \times 10^{-1}$ |
| PIPPack$_{RS}$ | $9.9999 \times 10^{-1}$ | $9.9997 \times 10^{-1}$ | $9.9999 \times 10^{-1}$ | $9.9999 \times 10^{-1}$ | $9.9999 \times 10^{-1}$ | $9.9999 \times 10^{-1}$ | $9.7988 \times 10^{-1}$ | $9.9611 \times 10^{-1}$ | $9.9999 \times 10^{-1}$ | $9.9999 \times 10^{-1}$ | $9.9999 \times 10^{-1}$ |
| PIPPack$_{ensembled}$ | $9.9999 \times 10^{-1}$ | $9.9993 \times 10^{-1}$ | $9.9999 \times 10^{-1}$ | $9.9999 \times 10^{-1}$ | $9.9999 \times 10^{-1}$ | $9.9975 \times 10^{-1}$ | $9.0331 \times 10^{-1}$ | $9.4465 \times 10^{-1}$ | $9.9999 \times 10^{-1}$ | $9.9999 \times 10^{-1}$ | $9.9999 \times 10^{-1}$ |
| PIPPack$_{ensembled + RS}$ | $9.9999 \times 10^{-1}$ | $9.9997 \times 10^{-1}$ | $9.9999 \times 10^{-1}$ | $9.9999 \times 10^{-1}$ | $9.9999 \times 10^{-1}$ | $9.9999 \times 10^{-1}$ | $9.9889 \times 10^{-1}$ | $9.9911 \times 10^{-1}$ | $9.9999 \times 10^{-1}$ | $9.9999 \times 10^{-1}$ | $9.9999 \times 10^{-1}$ |
| DiffPack | $9.9999 \times 10^{-1}$ | $9.9999 \times 10^{-1}$ | $9.9999 \times 10^{-1}$ | $9.9999 \times 10^{-1}$ | $9.9999 \times 10^{-1}$ | $9.9999 \times 10^{-1}$ | $9.9587 \times 10^{-1}$ | $9.9999 \times 10^{-1}$ | $9.9999 \times 10^{-1}$ | $9.9999 \times 10^{-1}$ | $9.9999 \times 10^{-1}$ |
| DiffPack$_{conf}$ | $9.9999 \times 10^{-1}$ | $9.9999 \times 10^{-1}$ | $9.9999 \times 10^{-1}$ | $9.9999 \times 10^{-1}$ | $9.9999 \times 10^{-1}$ | $9.9999 \times 10^{-1}$ | $9.4876 \times 10^{-1}$ | $9.9987 \times 10^{-1}$ | $9.9999 \times 10^{-1}$ | $9.9999 \times 10^{-1}$ | $9.9999 \times 10^{-1}$ |
| AttnPacker | $9.9624 \times 10^{-1}$ | $9.9924 \times 10^{-1}$ | $8.6122 \times 10^{-1}$ | $9.9999 \times 10^{-1}$ | $9.9999 \times 10^{-1}$ | $9.9998 \times 10^{-1}$ | $5.0228 \times 10^{-1}$ | $9.9999 \times 10^{-1}$ | $9.9999 \times 10^{-1}$ | $9.9999 \times 10^{-1}$ | $9.9999 \times 10^{-1}$ |
| AttnPacker$_{PP}$ | $9.9999 \times 10^{-1}$ | $9.9999 \times 10^{-1}$ | $9.9999 \times 10^{-1}$ | $9.9999 \times 10^{-1}$ | $9.9999 \times 10^{-1}$ | $9.9998 \times 10^{-1}$ | $8.0496 \times 10^{-1}$ | $9.9999 \times 10^{-1}$ | $9.9999 \times 10^{-1}$ | $6.6530 \times 10^{-1}$ | $9.9987 \times 10^{-1}$ |
| DLPacker$_{seq}$ | $9.9999 \times 10^{-1}$ | $9.9999 \times 10^{-1}$ | $9.9999 \times 10^{-1}$ | $9.9999 \times 10^{-1}$ | $9.9999 \times 10^{-1}$ | $9.9999 \times 10^{-1}$ | $9.9999 \times 10^{-1}$ | $9.9999 \times 10^{-1}$ | $9.9999 \times 10^{-1}$ | $9.9999 \times 10^{-1}$ | $9.9999 \times 10^{-1}$ |
| DLPacker$_{natoms}$ | $9.9999 \times 10^{-1}$ | $9.9999 \times 10^{-1}$ | $9.9999 \times 10^{-1}$ | $9.9999 \times 10^{-1}$ | $9.9999 \times 10^{-1}$ | $9.9999 \times 10^{-1}$ | $9.9999 \times 10^{-1}$ | $9.9999 \times 10^{-1}$ | $9.9999 \times 10^{-1}$ | $9.9999 \times 10^{-1}$ | $9.9999 \times 10^{-1}$ |
| DLPacker$_{score}$ | $9.9999 \times 10^{-1}$ | $9.9999 \times 10^{-1}$ | $9.9999 \times 10^{-1}$ | $9.9999 \times 10^{-1}$ | $9.9999 \times 10^{-1}$ | $9.9999 \times 10^{-1}$ | $9.9999 \times 10^{-1}$ | $9.9999 \times 10^{-1}$ | $9.9999 \times 10^{-1}$ | $9.9999 \times 10^{-1}$ | $9.9999 \times 10^{-1}$ |
| FASPR | $9.9999 \times 10^{-1}$ | $9.9999 \times 10^{-1}$ | $9.9999 \times 10^{-1}$ | $9.9999 \times 10^{-1}$ | $9.9999 \times 10^{-1}$ | $9.9999 \times 10^{-1}$ | $9.9940 \times 10^{-1}$ | $9.9999 \times 10^{-1}$ | $9.9999 \times 10^{-1}$ | $9.9999 \times 10^{-1}$ | $9.9999 \times 10^{-1}$ |
| PyRosetta | $9.9999 \times 10^{-1}$ | $9.9999 \times 10^{-1}$ | $9.9999 \times 10^{-1}$ | $9.9999 \times 10^{-1}$ | $9.9999 \times 10^{-1}$ | $9.9950 \times 10^{-1}$ | $9.9873 \times 10^{-1}$ | $9.9999 \times 10^{-1}$ | $9.9999 \times 10^{-1}$ | $9.9999 \times 10^{-1}$ | $9.9999 \times 10^{-1}$ |
| SCWRL4 | $9.9999 \times 10^{-1}$ | $9.9999 \times 10^{-1}$ | $9.9999 \times 10^{-1}$ | $9.9999 \times 10^{-1}$ | $9.9999 \times 10^{-1}$ | $9.9999 \times 10^{-1}$ | $9.9999 \times 10^{-1}$ | $9.9999 \times 10^{-1}$ | $9.9999 \times 10^{-1}$ | $9.9999 \times 10^{-1}$ | $9.9999 \times 10^{-1}$ |
| **Repacking AlphaFold3's Predicted Backbones** | | | | | | | | | | | |
| FlowPacker$_{BC40}$ | $9.9999 \times 10^{-1}$ | $9.8380 \times 10^{-1}$ | $9.9999 \times 10^{-1}$ | $9.9999 \times 10^{-1}$ | $9.9999 \times 10^{-1}$ | $9.7180 \times 10^{-1}$ | $9.9998 \times 10^{-1}$ | $9.9929 \times 10^{-1}$ | $9.9999 \times 10^{-1}$ | $9.9999 \times 10^{-1}$ | $9.9999 \times 10^{-1}$ |
| FlowPacker$_{BC40, conf}$ | $9.9999 \times 10^{-1}$ | $9.9939 \times 10^{-1}$ | $9.9999 \times 10^{-1}$ | $9.9999 \times 10^{-1}$ | $9.9999 \times 10^{-1}$ | $9.9620 \times 10^{-1}$ | $9.9950 \times 10^{-1}$ | $9.9966 \times 10^{-1}$ | $9.9999 \times 10^{-1}$ | $9.9999 \times 10^{-1}$ | $9.9999 \times 10^{-1}$ |
| FlowPacker$_{cluster}$ | $9.9998 \times 10^{-1}$ | $9.9068 \times 10^{-1}$ | $9.9997 \times 10^{-1}$ | $9.9989 \times 10^{-1}$ | $9.9987 \times 10^{-1}$ | $9.1634 \times 10^{-1}$ | $9.9443 \times 10^{-1}$ | $7.8500 \times 10^{-1}$ | $9.9999 \times 10^{-1}$ | $9.9999 \times 10^{-1}$ | $9.9998 \times 10^{-1}$ |
| FlowPacker$_{cluster, conf}$ | $9.9990 \times 10^{-1}$ | $9.9371 \times 10^{-1}$ | $9.9986 \times 10^{-1}$ | $9.9940 \times 10^{-1}$ | $9.9996 \times 10^{-1}$ | $9.9717 \times 10^{-1}$ | $9.8461 \times 10^{-1}$ | $7.7329 \times 10^{-1}$ | $9.9999 \times 10^{-1}$ | $9.9999 \times 10^{-1}$ | $9.9998 \times 10^{-1}$ |
| PIPPack | $9.9999 \times 10^{-1}$ | $9.9999 \times 10^{-1}$ | $9.9999 \times 10^{-1}$ | $9.9999 \times 10^{-1}$ | $9.9999 \times 10^{-1}$ | $9.8558 \times 10^{-1}$ | $9.9992 \times 10^{-1}$ | $9.9884 \times 10^{-1}$ | $9.9999 \times 10^{-1}$ | $9.9999 \times 10^{-1}$ | $9.9999 \times 10^{-1}$ |
| PIPPack$_{RS}$ | $9.9999 \times 10^{-1}$ | $9.9998 \times 10^{-1}$ | $9.9999 \times 10^{-1}$ | $9.9999 \times 10^{-1}$ | $9.9999 \times 10^{-1}$ | $9.9975 \times 10^{-1}$ | $9.9989 \times 10^{-1}$ | $9.9982 \times 10^{-1}$ | $9.9999 \times 10^{-1}$ | $9.9999 \times 10^{-1}$ | $9.9995 \times 10^{-1}$ |
| PIPPack$_{ensembled}$ | $9.9999 \times 10^{-1}$ | $9.9999 \times 10^{-1}$ | $9.9999 \times 10^{-1}$ | $9.9998 \times 10^{-1}$ | $9.9999 \times 10^{-1}$ | $9.0477 \times 10^{-1}$ | $9.9787 \times 10^{-1}$ | $9.9979 \times 10^{-1}$ | $9.9999 \times 10^{-1}$ | $9.9999 \times 10^{-1}$ | $9.9999 \times 10^{-1}$ |
| PIPPack$_{ensembled + RS}$ | $9.9999 \times 10^{-1}$ | $9.9999 \times 10^{-1}$ | $9.9999 \times 10^{-1}$ | $9.9999 \times 10^{-1}$ | $9.9997 \times 10^{-1}$ | $9.9975 \times 10^{-1}$ | $9.9963 \times 10^{-1}$ | $9.9999 \times 10^{-1}$ | $9.9999 \times 10^{-1}$ | $9.9999 \times 10^{-1}$ | $9.9992 \times 10^{-1}$ |
| DiffPack | $9.9999 \times 10^{-1}$ | $9.9999 \times 10^{-1}$ | $9.9999 \times 10^{-1}$ | $9.9999 \times 10^{-1}$ | $9.9999 \times 10^{-1}$ | $9.9999 \times 10^{-1}$ | $9.9999 \times 10^{-1}$ | $9.9999 \times 10^{-1}$ | $9.9999 \times 10^{-1}$ | $9.9999 \times 10^{-1}$ | $9.9999 \times 10^{-1}$ |
| DiffPack$_{conf}$ | $9.9999 \times 10^{-1}$ | $9.9999 \times 10^{-1}$ | $9.9999 \times 10^{-1}$ | $9.9999 \times 10^{-1}$ | $9.9999 \times 10^{-1}$ | $9.9999 \times 10^{-1}$ | $9.9999 \times 10^{-1}$ | $9.9999 \times 10^{-1}$ | $9.9999 \times 10^{-1}$ | $9.9999 \times 10^{-1}$ | $9.9999 \times 10^{-1}$ |
| AttnPacker | $9.8310 \times 10^{-1}$ | $9.9999 \times 10^{-1}$ | $4.1086 \times 10^{-1}$ | $9.9999 \times 10^{-1}$ | $9.9999 \times 10^{-1}$ | $9.9999 \times 10^{-1}$ | $9.8299 \times 10^{-1}$ | $9.9999 \times 10^{-1}$ | $9.9999 \times 10^{-1}$ | $9.9999 \times 10^{-1}$ | $9.9999 \times 10^{-1}$ |
| AttnPacker$_{PP}$ | $9.9999 \times 10^{-1}$ | $9.9999 \times 10^{-1}$ | $9.9998 \times 10^{-1}$ | $9.9999 \times 10^{-1}$ | $9.9999 \times 10^{-1}$ | $9.9999 \times 10^{-1}$ | $9.9487 \times 10^{-1}$ | $9.9999 \times 10^{-1}$ | $9.9999 \times 10^{-1}$ | $8.1172 \times 10^{-10}$* | $1.7432 \times 10^{-1}$ |
| DLPacker$_{seq}$ | $9.9999 \times 10^{-1}$ | $9.9999 \times 10^{-1}$ | $9.9999 \times 10^{-1}$ | $9.9999 \times 10^{-1}$ | $9.9999 \times 10^{-1}$ | $9.9999 \times 10^{-1}$ | $9.9999 \times 10^{-1}$ | $9.9999 \times 10^{-1}$ | $9.9999 \times 10^{-1}$ | $9.9999 \times 10^{-1}$ | $9.9999 \times 10^{-1}$ |
| DLPacker$_{natoms}$ | $9.9999 \times 10^{-1}$ | $9.9999 \times 10^{-1}$ | $9.9999 \times 10^{-1}$ | $9.9999 \times 10^{-1}$ | $9.9999 \times 10^{-1}$ | $9.9999 \times 10^{-1}$ | $9.9999 \times 10^{-1}$ | $9.9999 \times 10^{-1}$ | $9.9999 \times 10^{-1}$ | $9.9999 \times 10^{-1}$ | $9.9999 \times 10^{-1}$ |
| DLPacker$_{score}$ | $9.9999 \times 10^{-1}$ | $9.9999 \times 10^{-1}$ | $9.9999 \times 10^{-1}$ | $9.9999 \times 10^{-1}$ | $9.9999 \times 10^{-1}$ | $9.9999 \times 10^{-1}$ | $9.9999 \times 10^{-1}$ | $9.9999 \times 10^{-1}$ | $9.9999 \times 10^{-1}$ | $9.9999 \times 10^{-1}$ | $9.9999 \times 10^{-1}$ |
| FASPR | $9.9999 \times 10^{-1}$ | $9.9999 \times 10^{-1}$ | $9.9999 \times 10^{-1}$ | $9.9999 \times 10^{-1}$ | $9.9999 \times 10^{-1}$ | $9.9999 \times 10^{-1}$ | $9.9998 \times 10^{-1}$ | $9.9999 \times 10^{-1}$ | $9.9999 \times 10^{-1}$ | $9.9999 \times 10^{-1}$ | $9.9999 \times 10^{-1}$ |
| PyRosetta | $9.9999 \times 10^{-1}$ | $9.9999 \times 10^{-1}$ | $9.9999 \times 10^{-1}$ | $9.9999 \times 10^{-1}$ | $9.9999 \times 10^{-1}$ | $9.9999 \times 10^{-1}$ | $9.9968 \times 10^{-1}$ | $9.9999 \times 10^{-1}$ | $9.9999 \times 10^{-1}$ | $4.7213 \times 10^{-1}$ | $9.8044 \times 10^{-1}$ |
| SCWRL4 | $9.9999 \times 10^{-1}$ | $9.9999 \times 10^{-1}$ | $9.9999 \times 10^{-1}$ | $9.9999 \times 10^{-1}$ | $9.9999 \times 10^{-1}$ | $9.9997 \times 10^{-1}$ | $9.9999 \times 10^{-1}$ | $9.9999 \times 10^{-1}$ | $9.9999 \times 10^{-1}$ | $9.9999 \times 10^{-1}$ | $9.9999 \times 10^{-1}$ |

Note: Asterisks (*) indicate values that are statistically significantly better than the corresponding value from AlphaFold's side-chain predictions.

### 3 Details of Our Algorithm for Repacking AlphaFold Side-Chains Using a Backbone Confidence-Aware Integrative Approach

Our script to generate side-chains that are a combination of the other side-chain packing tools is primarily designed as a post-prediction correction step for AlphaFold's protein structure prediction pipeline. The full procedure, as summarized in the main paper, is given in **Algorithm 1**.

We refer to the number of amino acid residues in the given protein as $r$, and the maximum number of heavy atoms per residue for any given amino acid is 14 as specified in AlphaFold's internal full-atom model of protein tertiary structures (`https://github.com/google-deepmind/alphafold/blob/2ca41b1991bbf17be4a86c1face55597ba4c7621/alphafold/model/all_atom.py#L23`). Additionally, the backbones stay constant throughout the algorithm and are only included for correctness, as the scoring of the protein and writing of the final pose in PyRosetta involve the full protein structure. As such, the computation of each residue's backbone confidence score is done beforehand, and it divides the sum (across the backbone atoms' confidences) by 100 to scale down the plDDT values from a range of $[0, 100]$ to $[0, 1]$ and then takes the average across those 4 atoms.

**Algorithm 1** Backbone confidence-aware integrative approach for repacking AlphaFold side-chains

**Input:**

`pdb_path`: AlphaFold-outputted PDB file holding protein structure
`tools`: List of side-chain packing tools whose predictions to draw from
`output_pdb_path`: Path to save new PDB file to

**Output:**

Protein structure with updated side-chains is saved to `output_pdb_path`

**function** RefineSideChains(`pdb_path`, `tools`, `output_pdb_path`)
    ▷ *Retrieves all $\chi$ angles needed*
    $\mathbf{B}_{i,j}$ := GetBackbones(`pdb_path`) ▷ Backbone atoms $\in \mathbb{R}^{r\times 14}$ remain constant
    $\boldsymbol{\chi}_{i,j}$ := GetDihedrals(`pdb_path`) ▷ $\boldsymbol{\chi} \in \mathbb{R}^{r\times 4}$
    $\mathbf{p}_{i,j}$ := GetAtomConfidences(`pdb_path`) ▷ plDDT values $\in \mathbb{R}^{r\times 14}$
    **for each** `tool` **in** `tools` **do**
        $\boldsymbol{\chi}^{\texttt{tool}}_{i,j}$ := RepackSideChains(`tool`, `pdb_path`) ▷ $\boldsymbol{\chi}^{\texttt{tool}} \in \mathbb{R}^{r\times 4}$
        `alternate_chis[tool]` := $\boldsymbol{\chi}^{\texttt{tool}}_{i,j}$
    **end for**

    ▷ *Computes backbone confidences*
    **for each** $i$ **in** $\{1, 2, \dots, r\}$ **do**
        $\mathbf{c}_i := \frac{\sum_{\text{atom}\in\{N, C_\alpha, C, O\}} \mathbf{p}_{i,\text{atom}}}{400}$ ▷ $\mathbf{c} \in \mathbb{R}^r$
    **end for**

    ▷ *Runs greedy optimization*
    `score_fxn` := InitializeScoreFunction(REF15)
    `curr_score` := `score_fxn`($\mathbf{B}$, $\boldsymbol{\chi}$)
    **for each** `iter` **in** $\{1, 2, \dots,$ `num_iters`$\}$ **do**
        `res` := RandomInteger(`low=`1, `high=`$r$)
        `chi_num` := RandomInteger(`low=`1, `high=`min(4, NumChis($\boldsymbol{\chi}$, `residue=res`)))
        `tool` := RandomChoice(`tools`)

        `old_chi` := $\boldsymbol{\chi}_{\texttt{res},\texttt{chi\_num}}$
        `new_chi` := $\mathbf{c}_{\texttt{res}}$ · `old_chi` + $(1 - \mathbf{c}_{\texttt{res}})$ · `alternate_chis[tool]`$_{\texttt{res},\texttt{chi\_num}}$
        $\boldsymbol{\chi}_{\texttt{res},\texttt{chi\_num}}$ := `new_chi`
        `new_score` := `score_fxn`($\mathbf{B}$, $\boldsymbol{\chi}$)

        **if** `new_score` < `curr_score` **then**
            `curr_score` := `new_score`
        **else**
            $\boldsymbol{\chi}_{\texttt{res},\texttt{chi\_num}}$ := `old_chi`
        **end if**
    **end for**

    WritePDB($\mathbf{B}$, $\boldsymbol{\chi}$, `output_pdb_path`)
**end function**
